# The genetic requirements for HiVir-mediated onion necrosis by *Pantoea ananatis*, a necrotrophic plant pathogen

**DOI:** 10.1101/2022.11.22.517531

**Authors:** Gi Yoon Shin, Bhabesh Dutta, Brian Kvitko

**Author notes:** Corresponding author: Brian Kvitko.

## Abstract

*Pantoea ananatis* is an unusual bacterial pathogen that lacks typical virulence determinants yet causes extensive necrosis in onion foliage and bulb tissues. The onion necrosis phenotype is dependent on the expression of a phosphonate toxin, pantaphos that is catalyzed by putative enzymes encoded by the HiVir gene cluster. The genetic contributions of individual *hvr* genes in HiVir-mediated onion necrosis remain largely unknown except for the first gene *hvrA* (phosphoenolpyruvate mutase, *pepM*) whose deletion resulted in the loss of onion pathogenicity. In this study, using gene deletion mutation and complementation, we report that of the ten remaining genes, *hvrB*-*hvrF* are also strictly required for the HiVir-mediated onion necrosis and *in planta* bacterial growth whereas *hvrG*-*hvrJ* partially contributed to these phenotypes. As the HiVir gene cluster is a common genetic feature shared among the onion-pathogenic *P. ananatis* strains, and as it could serve as a useful diagnostic marker of onion pathogenicity, we sought to understand the genetic basis of HiVir positive yet phenotypically deviant (non-pathogenic) strains. We identified and genetically characterized inactivating single nucleotide polymorphisms (SNPs) in essential *hvr* genes of six phenotypically deviant *P. ananatis* strains. Finally, inoculation of the cell-free spent medium of P_tac_-driven HiVir strain caused *P. ananatis*-characteristic red onion scale necrosis (RSN) as well as cell death symptoms in tobacco. The co-inoculation of the spent medium with essential *hvr* mutant strains restored strains’ *in planta* populations to the wild-type level, suggesting that necrosis is important for proliferation of *P. ananatis* in onion tissue.

## Introduction

Onion Center Rot caused by the gram-negative, bacterial pathogen *Pantoea ananatis* is an economically important disease. The infection of onion leaves leads to the development of water-soaked lesions that turn necrotic and extend down to the neck of the onion bulb (Gitaitis and Gay, 1997). Through the infected neck, the pathogen gains an entry into the bulb where it causes brown discoloration and necrosis of the internal scales, resulting in significant economic losses in both pre- and post-harvest conditions (Carr et al. 2013). Recent efforts were largely focused on understanding virulence mechanism of *Pantoea* spp. which tend to be unique compared to other members in Erwiniaceae family (Asselin et al. 2018; Takikawa et al. 2018; Stice et al., 2020; Polidore et al. 2021). *P. ananatis* is an atypical gram negative plant pathogen in that it neither possesses a type II secretion system for the deployment of plant cell wall degrading enzymes nor type III secretion systems for translocation of effectors into plant cells to disarm the plant immunity (De Maayer et al. 2014). Instead, onion pathogenicity by this necrotrophic pathogen requires the HiVir (<u>Hi</u>gh <u>Vir</u>ulence) biosynthetic cluster to produce phosphonate toxin called pantaphos. In 2018, Asselin *et al*. discovered the HiVir gene cluster, determined that it was a common feature of onion pathogenic *P. ananatis* strains and predicted that it might encode a biosynthetic pathway for a phosphonate compound. The HiVir containing strains caused necrosis on onion leaves, bulbs, and scales (Asselin et al. 2018; Stice et al. 2020) as well as HR-like cell death in tobacco in a manner that is dependent on the *hvrA*/*pepM* gene for phosphoenolpyruvate mutase, the typical first committed enzymatic step in the synthesis of phosphonates. Polidore et al. (2021) identified and purified 2-(hydroxy[phosphono]methyl)maleate or ‘pantaphos’ as the phosphonate compound produced in by *P. ananatis* in a HiVir-dependent manner. Pantaphos was comparably phytotoxic to the phosphonate herbicide glyphosate in a radish seedling assay and the injection of the purified pantaphos into onion bulbs resulted in extensive bulb necrosis.

The HiVir gene cluster contains eleven genes (designated as *hvr* genes by Polidore et al.) that are arranged in a manner consistent with expression as a single operon. According to the pantaphos biosynthetic pathway proposed by Polidore et al., six of eleven *hvr* genes are involved in the production of pantaphos (Polidore et al. 2021). Namely these are *hvrA* (phosphoenolpyruvate mutase, PepM), *hvrB* (flavin-dependent monooxygenase), *hvrC* (phosphonomethylmalate synthase), *hvrD* (isopropylmalate dehydratase-like protein, large subunit), *hvrE* (isopropylemalate dehydratase-like protein) and *hvrK* (flavin reductase). In addition, the *hvrI*-encoded MFS transporter was predicted to be employed for pantaphos export (Asselin et al. 2018; Polidore et al. 2021). Of the 11 *hvr* genes, only two have been tested independently for their genetic contributions to onion necrosis. These genes were *hvrA* (phosphoenolpyruvate mutase, PepM), which is required for all onion necrosis phenotypes (Asselin et al. 2018; Stice et al. 2021) and *hvrI* (MFS transporter protein), which resulted in reduced foliar necrosis when mutated (Asselin et al. 2018).

The HiVir gene cluster has been a useful genetic feature to identify onion-pathogenic *P. ananatis* strains (Asselin et al. 2018; Stice et al. 2020; Agarwal et al. 2021). However, Polidore et al. (2021), and Agarwal et al (2021) identified several *P. ananatis* strains that encode the HiVir cluster but lack the capacity to induce necrosis in onions (Agarwal et al. 2021; Polidore et al. 2021). We hypothesized that these phenotypically deviant strains might carry inactivating SNPs in *hvr* genes required for necrosis.

In this study, we generated in-frame deletions of *hvr* genes in *P. ananatis* PNA 97-1R to determine their individual genetic contributions to onion scale necrosis, *in planta* bacterial population and the production of onion foliar lesions. We determined that *hvrA-hvrF* were required for necrosis while *hvrG-hvrJ* contributed to necrosis-associated phenotypes but were not strictly required. These results largely but incompletely overlap with predicted requirements from the proposed pantaphos biosynthetic pathway. Furthermore, we genetically characterized inactivating SNPs in necrosis-required *hvr* genes for six phenotypically deviant (HiVir+ RSN-) *P. ananatis* strains.

## Method

### Bacterial growth conditions

Bacterial strains used in this study are listed in Table S1. *P. ananatis* and *E. coli* DH5α, MaH1, and RHO5 strains were routinely grown overnight in/on Lysogeny Broth (LB: 10 g/L tryptone, 5 g/L yeast extract, 5 g/L NaCl) broth or agar (15 g/L agar) at 28 ºC and at 37 ºC, respectively. The growth medium was supplemented with antibiotics at the following final concentrations; chloramphenicol 50 µg/ml, gentamicin 15 µg/ml, kanamycin 50 µg/ml, rifampicin 40 µg/ml, tetracycline 15 µg/ml and trimethoprim 40 µg/ml as appropriate.

### Plant growth conditions

Onion seedlings (cv. Century) were potted in 16 cm X 15 cm (diameter X height) plastic pots with SunGrow 3B commercial potting mix on December 13, 2021. The onions were grown in the greenhouse until the third week of March 2022 when they were inoculated for foliar tip assays.

### Creation of *hvr* gene deletion mutants and HiVir-inducible *P. ananatis*

To determine the genetic basis of HiVir-mediated onion pathogenicity, individual *hvr* gene deletion mutant strains of *P. ananatis* PNA 97-1R were generated by allelic exchange. To avoid disruption of the expression of up- and downs-tream *hvr* genes, up to seven codons from the start and the end of the target *hvr* open reading frame were included in the 450 bp up- and down-stream flanks. In the middle of the two flanks, a restriction site (SmaI) was introduced and, *attB* sequences were attached at the either ends of the merged flanks to make it Gateway BP cloning compatible. The complexity of the deletion construct was assessed by the gBlocks® Gene Fragments Entry tool at IDT website (https://eu.idtdna.com/site/order/gblockentry). Subsequently, approved fragments were synthesized by TWIST BIOSCIENCE (San Francisco, CA). The sequences of the deletion fragments are listed in Table S2.

The insertion of deletion fragment into the vector pR6KT2G (Stice et al. 2020) was conducted using Gateway™ BP Clonase™ II Enzyme Mix (Invitrogen) following the manufacturer’s protocol. A total volume of inactivated BP reaction was placed on the VMWP membrane (Millipore) suspended in sterile dH_2_0 for 30 min to remove excess salt before being electroporated into *E. coli* MaH1 (Kvitko et al., 2012). *E. coli* MaH1 transformants were selected on LB agar amended with gentamicin after overnight incubation. To verify that BP reaction had successfully taken place, a putative recombinant vector was digested and confirmed using SmaI, and Sanger sequenced using a pR6KT2G-specific primer pair (Table S3) with the following PCR reaction conditions: 12.5 µl of GoTaq® Green Master Mix (Promega), 1 µl of each primer at 10 µM concentration, 1 µl of 50 ng/µl DNA, and 9.5 µl of nuclease-free water to make up a total of 25 µl PCR reaction mixture. The PCR cycling conditions were as follows: initial denaturation at 95 ºC for 5 min, 30 cycles of 30 sec denaturation at 95 ºC, 30 sec of annealing at 60 ºC (Table S3), 1 min/kb extension at 72 ºC, then 5 min of final extension at 72 ºC before holding at 4 ºC indefinitely. The PCR amplicon was visualized in 1 % agarose gel stained with SYBR safe DNA gel stain (Invitrogen), cleaned with Monarch PCR & DNA Cleanup Kit (NEB), and sequenced at Eurofins Genomics USA. Subsequent steps leading to the allelic exchange-mediated *hvr* gene deletion mutations in *P. ananatis* PNA 97-1R were carried out by following the method described by Stice et al. (2020). The putative deletion mutants were screened and confirmed by colony PCR and sequencing using “Out” primers (Table S3).

To insert IPTG (isopropylthio-β-galactoside) inducible promoter (P_tac_) in the upstream of chromosomal *hvrA* gene of *P. ananatis* PNA 97-1R, pSNARE was created from Pre-SNARE and pSC201. The Pre-SNARE fragment that carries *lacIq* gene, IPTG inducible promoter (Ptac), optimized ribosome binding site (RBS) and 450 bp partial *hvrA* gene (Table S2) was synthesized by TWIST BIOSCIENCE (San Francisco, CA). The native RBS of *hvrA* was customized for the optimal translation initiation using UTR designer (https://sbi.postech.ac.kr/utr_designer) and RBS calculator v 2.1 (https://salislab.net/software/predict_rbs_calculator). The Pre-SNARE was PCR amplified using Pre-SNARE_SphI_F and Pre-SNARE_NsiI_R primers (Table S3), restricted and ligated with NsiI and SphI digested pSC201 using Gibson Assembly® Master Mix (NEB) following the manufacturer’s protocol. The modification of pSC201 into pSNARE was confirmed by XhoI and XbaI double restriction and sequencing using lacIQ_F and test_pSC201_R primers (Table S3). The single cross-over between pSNARE and the chromosome of *P. ananatis* PNA97-1R was facilitated by biparental mating as described by Stice et al. (2020), which was maintained in the host genome by selecting the host cell’s growth with trimethoprim. The insertion was validated by PCR and sequencing using primers lacIQ_F and hvrA_R (Table S3).

### Creation of *hvr* gene deletion complementing strains

The complementation of *hvr* mutation was achieved by PCR amplifying the full copy of an individual *hvr* gene as well as up to 40 bp upstream sequence to include the native RBS sequence using “Comp” primers (Table S3). The PCR product was gel-purified using Monarch DNA Gel Extraction Kit (NEB) and BP cloned into the entry vector pDONR221 as described above. The transformed *E. coli* DH5α colonies were recovered on LB agar amended with kanamycin and the presence of the recombinant pDONR221 vector was confirmed by the colony PCR using M13 primers (Table S3) and chloramphenicol sensitivity. The recombinant pDONR221 was extracted using GeneJet Plasmid MinPrep Kit (ThermoScientific), which was then cloned into pBS46 by LR reaction using Gateway™ LR Clonase™ II Enzyme Mix (Invitrogen). Finally, construction of *hvr* complementing pBS46 plasmid was confirmed by sequencing with an M13 primer followed by the electroporation of complementing plasmid into each corresponding electrocompetent *hvrB, hvrC, hvrD, hvrE, hvrF, hvrG, hvrH, hvrI, hvrJ* and *hvrK* deletion mutant strains of *P. ananatis* PNA 97-1R.

### Bacterial inoculum preparation

The *hvr* mutant and complementing strains of *P. ananatis* grown overnight in LB supplemented with antibiotics as appropriate, were pelleted and resuspended in sterile 1X phosphate buffered saline solution to 0.3 optical densities (OD_600_), which correlates with approximately 1×10^8^ CFU/ml. For foliar and scale necrosis assay, 10 µl of 1×10^8^ CFU/ml inoculum suspension (10^6^ CFU) was inoculated. For *in planta* bacterial population quantification in red onion scales, bacterial suspension was firstly standardized to OD_600nm_ = 0.1 from which a ten-fold dilution was made (1×10^6^ CFU/ml). A volume of 10 µl of this dilution (correlates with 1×10^4^ CFU) was used. For colony-stab inoculation, *hvr* mutant and complementing strains were grown on LB agar amended with antibiotics as appropriate, for 48 hours. The CFU of a single colony was calculated by performing a serial dilution which correlated with approximately 1×10^10^ CFU per colony.

### Red onion scale necrosis (RSN) assay

Red onion scale necrosis assay, previously described by Stice et al. (2018, 2020) was conducted with slight modifications. Commercially available red onion bulbs were purchased and sliced carefully to create approximately 3 cm X 3 cm unmarred scales. The scales were surface sterilized in 3 % sodium hypochlorite solution for 1 min thereafter rinsed twice in dH_2_0. Washed scales were placed on paper towels to air dry before placing them on autoclaved pipette tip trays. A few layers of paper towels were lined inside the clean plastic potting tray (27 cm X 52 cm) in which the towels were moistened with sterile dH_2_0. The pipette tip trays were placed on top of the wet towels and scales were positioned to not touch each other. A sterile 200 µl pipette tip was used to pierce the onion scales in the center where inoculum was deposited. Five scales per strain was inoculated with 10 µl of bacterial suspension (1×10^4^ CFU or 1×10^6^ CFU) as prepared described above. For colony stab inoculation, a single colony (1×10^10^ CFU) was scraped from a 48-hour old culture agar plate using a sterile 200 µl pipette tip, which was placed and left in the pierced onion scales for 4 days. Sterile saline solution was used as a negative control. After inoculation, the potting tray was covered with a second tray to create a “moist chamber” and incubated at room temperature for 4 days. This assay was repeated three times in total for data collection.

For the quantification of *in planta* bacterial population in onion scales, onion tissues at inoculation point were sampled at 4 days-post inoculation (dpi), using a metal borer (r = 2.5 mm), weighed, and placed in a 2 ml plastic tube filled with three 3mm zirconia beads (GlenMills^®^ Grinding Media) suspended in 500 µl of 1X phosphate buffered saline solution. The tissue was macerated using a SpeedMill PlUS homogenizer (Analytik Jena) for 2 min. The macerate was serially diluted to 10^−7^ in a sterile saline solution and 10 µl of diluents was plated out onto LB amended with Rifampicin and Gentamicin for the selection of mutant and complementing strains, respectively. After overnight incubation, visible colonies were counted, and CFU/gram of onion tissue was calculated for each scale sampled. Three scales were sampled per strain and this experiment was conducted at least three times.

### Onion foliar necrosis assay

The leaf tip assay was conducted on onion plants (cv. Century) established with at least 4 leaf blades in the greenhouse. The method described by Koirala et al. (2021) was followed with minor modifications. Three plants per bacterial strain were used as biological replicates and sterile 1X phosphate buffered saline solution was used as a negative control. Using surface sterilized pair of scissors, leaves were cut 1 cm from the apex and 10 µl of OD_600_= 0.3 bacterial suspension (1×10^6^ CFU) was deposited carefully onto the cut end. The lesion length was measured at 4 dpi. The onion foliar necrosis assay was repeated at least twice.

### Tobacco infiltration assay

The overnight culture of wildtype *P. ananatis* PNA 97-1R, HiVir-inducible strain (PNA 97-1R-SNARE::P_tac_*hvr*) and HiVir-inducible, *hvrF* deletion mutant strain (PNA 97-1R Δ*hvrF* -SNARE::P_tac_*hvr*) were grown in LM broth (10 g/L tryptone, 6 g/L yeast extract, 1.193 g/L KH_2_PO_4_, 0.6 g/L NaCl, 0.4 g/L MgSO_4_*7H_2_0 and 18 g/L agar) amended with trimethoprim as needed. After 24 h, a volume of overnight culture (50 µl) was transferred to fresh 5 ml of LM broth. The strains were grown in two conditions: IPTG (isopropylthio-β-galactoside) induced and non-IPTG induced. Under induced condition, 100 mM IPTG was added to the culture at a final concentration of 1 mM when the cell density has reached OD_600_ = 0.5 whereas no IPTG was added to the non-induced condition. After overnight incubation with shaking at 200 rpm, the cultures were centrifuged at 4 ºC for 10 min at 2585 rcf (eppendorf 5810 R). The supernatant of each culture was sterilized through a 0.2 µM filter to remove bacterial cells. Prior to tobacco infiltration, 10 µl of each filtrate was dropped on LB agar for 48 h incubation to ensure that filtrate was free of any bacterial contamination. After confirmation, approximately 100 µl of filtered spent media were syringe-infiltrated into tobacco (cv. xanthi SX) leaf panels under plant room conditions (12 h-day, 12 h-night, room temperature), thereafter, symptoms were observed for 3 days. LM broth with IPTG and no IPTG was used as a medium negative control.

### Co-inoculation assay using bacterial suspension and spent media

Inoculum suspension of wildtype (WT), Δ*hvrA*, Δ*hvrD* and *hvrA*_L216_ *P. ananatis* strains were prepared for the *in planta* population count as described above. An inoculum volume of 10 µl at 1×10^6^ CFU/ml (1×10^4^ CFU) was mixed with 10 µl of the cell-free HiVir-induced toxin <u>s</u>pent <u>m</u>edium (**tsm**) of HiVir-inducible (PNA 97-1R-SNARE::P_tac_*hvr*) and *hvrF* <u>m</u>utant <u>s</u>pent <u>m</u>edium (**msm**) of HiVir-inducible Δ*hvrF* deletion mutant (Δ*hvrF*-SNARE::P_tac_*hvr*). Controls included spent medium only, bacterial suspension only and sterile water. The RSN assay was performed as described above and scales were harvested at 4 dpi for *in planta* bacterial load count. This experiment was repeated twice.

### Statistical analysis

Statistical analyses and construction of the box plots were performed in RStudio (v 4.1.2) available at https://www.rstudio.com/. The normality of the dataset was determined using Bartlett test. For parametric data, ANOVA (Analysis of Variance) test was used and for non-parametric data, Kruskal-Wallis test was performed. The normalization of foliar lesion length data by 0.1 was carried out prior to statistical analysis.

### Identification of single nucleotide polymorphisms (SNPs) present in the *hvr* genes

The HiVir gene cluster was extracted from the genomes of 60 *P. ananatis* strains (*n*= 54 RSN^+^ / *n*= 6 RSN^-^) in which the HiVir gene cluster has been previously identified (Agarwal *et al*. 2021, Stice *et al*. 2021). This was carried out by BLASTn (Geneious Prime® 2022.2.2) search of *P. ananatis* LMG 2665^T^ HiVir gene cluster against the collection of genomes. Retrieved sequences were aligned using MAFFT (Geneious Prime® 2022.2.2) together with the HiVir sequence of a type strain *P. ananatis* LMG 2665^T^. Subsequently, all SNPs that resulted in the missense mutations as well as indels (insertions and deletions) in the coding sequence of HiVir gene cluster and that were not present in the reference/type HiVir sequence were manually recorded. The presence of SNPs in RSN^-^*P. ananatis* strains were further verified by sequencing the regions where the mutations have been identified. For the sequencing of *hvrB, hvrC* and *hvrF* genes, ‘Comp’ primers were used whereas newly designed ‘sequencing primers’ were employed for the sequencing of *hvrA* and *hvrI* genes (Table S3).

### Phylogenetic tree construction

The nucleotide sequence of *P. ananatis* PNA 97-1 HiVir gene cluster was used in the BLASTn search at NCBI. The homologous sequences of HiVir gene cluster were downloaded from BLAST search and aligned with *P. ananatis* PNA 97-1 HiVir gene cluster using MAFFT (Geneious Prime® 2022.2.2). Determination of the best-fit evolutionary model and construction of the maximum-likelihood phylogenetic tree were conducted using PhyML 3.0 (http://www.atgc-montpellier.fr/phyml/) and MEGA X (Kumar et al. 2018) was used to edit the tree.

### Creation of allelic exchange mutants and double *hvr* gene complementing strains

To investigate the impact of SNPs (D154G, L216S, I260K) in the HiVir-mediated RSN ability of the deviant (HiVir^+^ RSN^-^) *P. ananatis* strains (PANS 99-32, PANS 04-2, PANS 02-1, PNA 98-11, PNA 07-14, PNA 07-13), *hvrA* gene exchange experiment between the wildtype (HiVir^+^ RSN^+^ PNA 97-1R) and deviant *P. ananatis* strains (PANS 99-32, PANS 04-2, PANS 02-1, PNA 98-11, PNA 07-14, PNA 07-13) was set up. Firstly, different versions of *hvrA (hvrA*_*WT*_, *hvrA*_*D154G*_, *hvrA*_*L216S*_, *hvrA*_*I260K*_) gene were PCR amplified by the primers hvrA_exc_F and hvrA_exc_R that consist of *hvrA* gene specific sequences and *attB* sites (Table S3). Using the same method as described as above (See Creation of *hvr* gene deletion mutants), *hvrA* amplicon was BP-cloned into the allelic exchange vector pR6KT2G which was integrated into the chromosome of the target *P. ananatis* strain. The vector was cured via sucrose counter selection and the mutants were screened by RSN phenotyping and sequencing with hvrA_F and/or hvrA_R primer, (Table S3). Similarly, the wildtype copy of *hvrB*_WT_ was replaced with the *hvrB* gene of PANS 04-2 (*hvrB*_V91L_) in the PNA 97-1R genetic background. The effect of this missense mutation was evaluated by RSN ability of the *hvrB*_V91L_ carrying wildtype strain.

Additionally, pDONR221::*hvrC* was LR-cloned into pBAV226 (Vinatzer et al. 2006). The recombinant pBAV226 (pBAV226::*hvrC*) was verified using XbaI and XhoI double digest and was transformed into pCPP1383::*hvrA* complemented strains PANS 04-2 and PNA 07-13. The transformation was screened by tetracycline resistance and the complementation was confirmed by RSN phenotyping.

## Results

### Identification of *hvr* genes essential for HiVir-mediated red onion scale necrosis (RSN)

In continuation of the previous effort to identify genetic contribution of *hvr* gene in onion necrosis (Asselin et al. 2018), we created 10 *hvr* gene (*hvrB* to *hvrK*) deletion mutant strains of *P. ananatis* PNA 97-1R. These mutants were inoculated into red onion scales at different concentrations to determine residual necrosis resulting from low to high inoculum concentrations. We identified a set of *hvr* genes that were strictly required for the onion necrosis at all conditions well as the genes that were partially contributing to the phenotype in a dosage dependent manner. At 1×10^4^ CFU inoculum concentration, *hvrB-G* and *hvrJ* deletion mutant strains failed to cause red onion scale necrosis (RSN). The onion scales inoculated with Δ*hvrH* developed a halo around the inoculation point and necrosis was observed in scales inoculated with Δ*hvrI* and Δ*hvrK* (Figure 1A: Low). The RSN-mutants exhibited significantly lower *in planta* populations than that of the wildtype (WT) whereas WT comparable bacterial populations were recovered from the scales inoculated with Δ*hvrH*, Δ*hvrI* and Δ*hvrK* strains (Figure 1B). At 1×10^6^ CFU inoculum concentration, necrotic lesions formed in response to Δ*hvrH* and larger lesions were seen in Δ*hvrI* inoculated scales. At 1×10^10^CFU, RSN developed in Δ*hvrG* and Δ*hvrJ* infected scales when no necrosis was formed using 1×10^4^ to 1×10^6^ CFU (Figure 1A: High). For all three inoculum concentrations, Δ*hvrB*, Δ*hvrC*, Δ*hvrD*, Δ*hvrE* and Δ*hvrF* remained RSN-(Figure 1A). Thus, these genes were designated as essential genes for the HiVir-mediated RSN regardless of the inoculum concentration. The rest of *hvr* genes (*hvrG* to *hvrJ*) were assigned as partially contributing (to RSN) genes. However, *hvrK* gene was found to be non-essential for the HiVir-mediated RSN.

**Figure 1.**
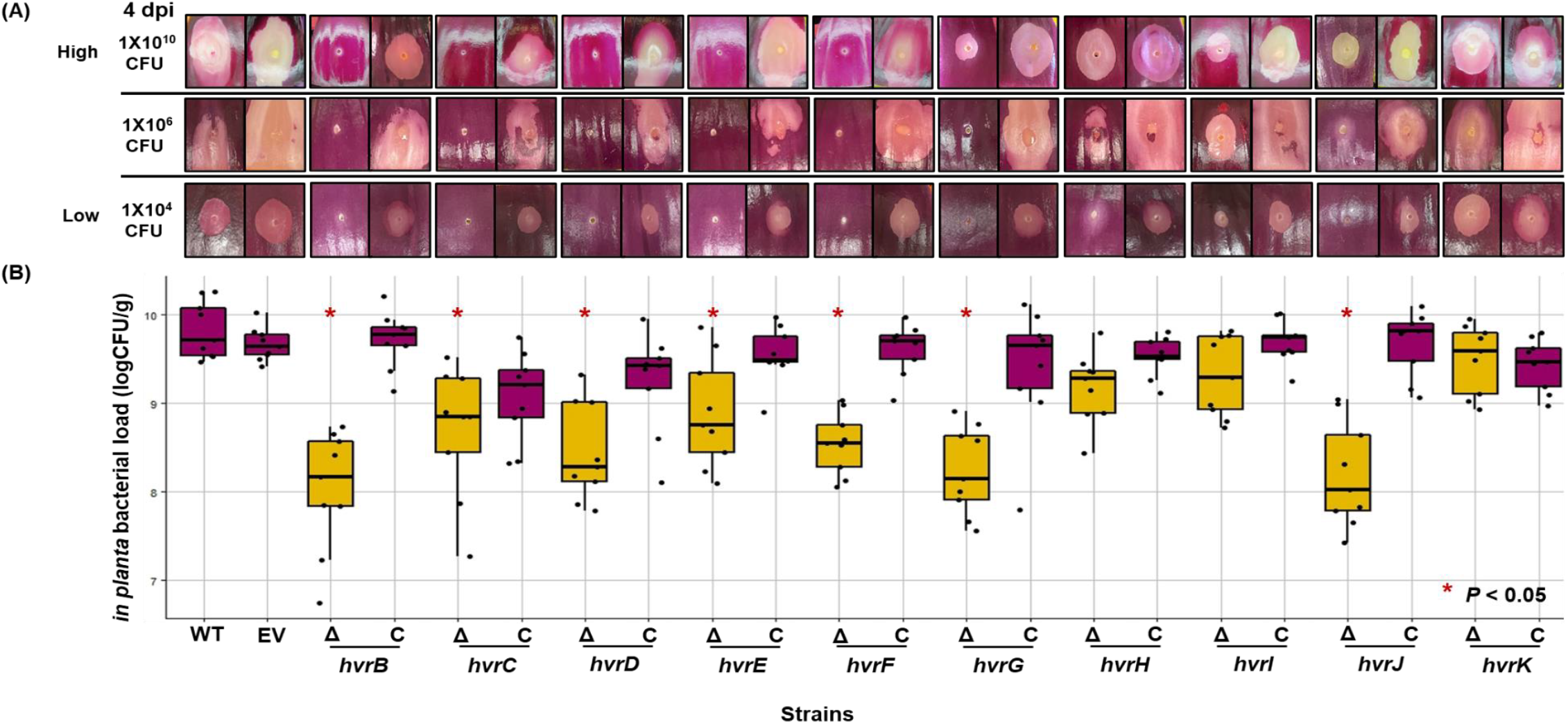
Red onion scale necrosis assay (RSN) and corresponding *in planta* population quantification of *hvr* gene deletion mutant and complementing strains of *Pantoea ananatis* PNA 97-1R. **(A) Top panel:** Red onion scales stab-inoculated with a colony (1×10^10^ CFU) at 4 days post inoculation (dpi), **Middle panel:** red onion scales inoculated with 1×10^6^ CFU at 4 dpi and, **(A) Bottom panel:** red onion scales inoculated with 1×10^4^ CFU at 4 dpi. The magenta-colored arrows represent essential *hvr* genes in HiVir-dependent RSN, pink = partially contributing and white = non-essential. **(B)** A box plot showing *in planta* bacterial population count per gram (logCFU/g) of onion scale tissue of onion sampled at 4 dpi. WT= wildtype PNA 97-1R, EV = wildtype strain carrying empty vector pBS46, Δ = *hvr* gene deletion mutant and C = *hvr* gene deletion complementing strain. *in planta* bacterial load of *hvr* gene deletion mutants was compared to that of the WT and significant reduction (*P* < 0.05) is indicated by the asterisk (*). Each jitter point represents an average count of a biological replicate. This experiment was repeated three times.

### Deletion of RSN essential *hvr* genes results in the loss of onion foliar necrosis

Consistent with the findings of the RSN assay, essential *hvr* gene mutants namely, Δ*hvrB*, Δ*hvrC*, Δ*hvrD*, Δ*hvrE* and Δ*hvrF* were unable to cause necrotic foliar lesions (Figure 2A). In addition to the essential genes, onion leaves inoculated with Δ*hvrJ* did not display any foliar necrosis at 4 dpi. Significantly reduced lesion sizes were observed in the leaves inoculated with Δ*hvrG*, Δ*hvrH* and Δ*hvrI* while Δ*hvrK* caused foliar lesion comparable to that of the wildtype (Figure 2B).

**Figure 2.**
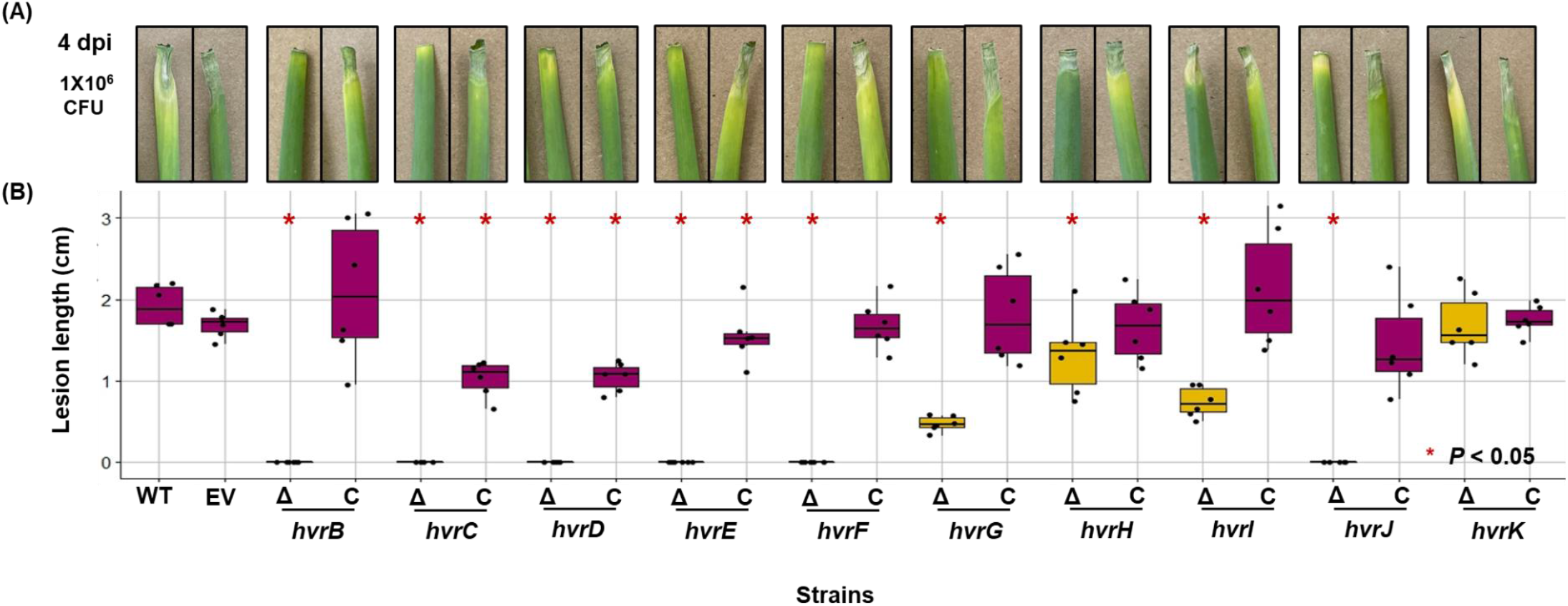
Onion leaf-tip assay. **(A)** Representative images of onion leaves inoculated with 1×10^6^ CFU of wildtype strain of *Pantoea ananatis* PNA 97-1R (WT), wildtype with empty vector pBS46 (EV), *hvr* gene deletion mutant (Δ) and complementing strains (C). The images were taken at 4 days post inoculation (dpi). **(B)** A box plot showing the lesion length of onion leaves inoculated with WT, EV, Δ and C strains at 4 dpi. The lesion length of *hvr* gene deletion mutants was compared to that of the WT and significant reduction in lesion length (*P* < 0.05) is indicated by the asterisk (*). Each jitter point represents a biological replicate. This experiment was repeated two times.

### HiVir+ RSN-deviant *P. ananatis* strains have SNPs in the open reading frame of essential *hvr* genes

To understand the genetic basis of RSN-phenotype in the HiVir+ deviant *P. ananatis* strains (PANS 99-32, PANS 04-2, PANS 02-1, PNA 98-11, PNA 07-14 and PNA 07-13), the nucleotide sequence of HiVir gene cluster was extracted from 54 HiVir+ *P. ananatis* strains (Table 1) and compared with the six deviant strains. Upon close inspection of the collective HiVir gene cluster at nucleotide and amino acid levels, we discovered unique SNPs and insertions were present in the essential *hvr* genes of the deviant strains that were not observed in the type strain *P. ananatis* LMG 2665^T^ and in other RSN^+^ strains. When translated, these SNPs gave rise to mis- and non-sense mutations as well as frameshift mutations, resulting in premature termination mediated by the stop codons. We identified missense mutations in the *hvrA* gene (phosphoenolpyruvate mutase, PepM) for 6 of 7 deviant strains representing three unique missense mutations (D154G, L216S, I260K) (Table 1). Other missense mutations included V98L change in *hvrB* (flavin-dependent monooxygenase) gene of PANS 04-2 and V116I change in *hvrI* (MFS transporter protein) gene of PANS 99-32. The insertion of adenine (4+A) and thymine (32+T) resulted in the frameshift of *hvrC* (phosphonomethylmalate synthase) open reading frame belonging to the strains PANS 04-2 and PNA 07-13, respectively. A nonsense mutation was found in the *hvrF* (O-methyltransferase) gene of PANS 99-32 at amino acid position 70.

**Table 1.**
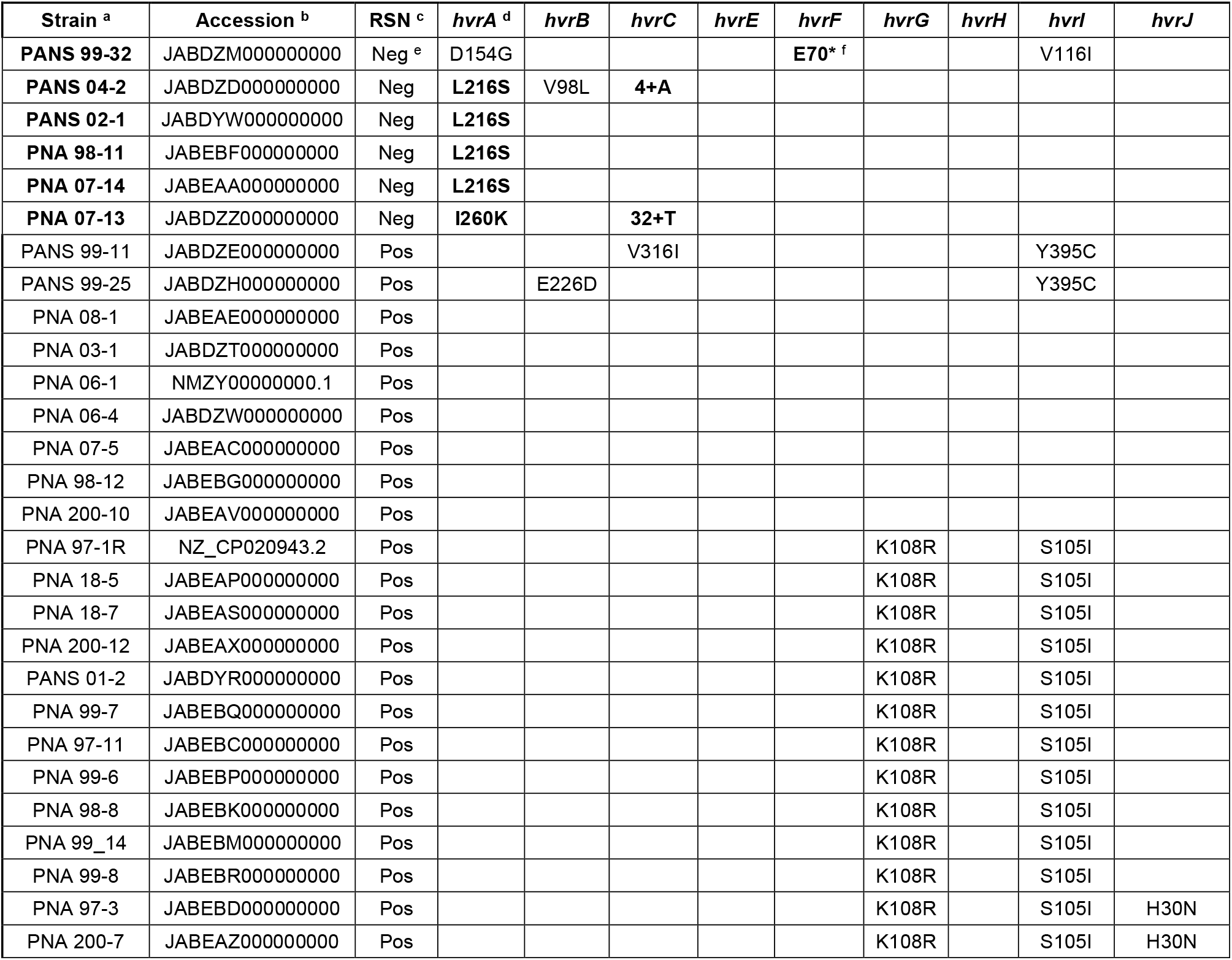
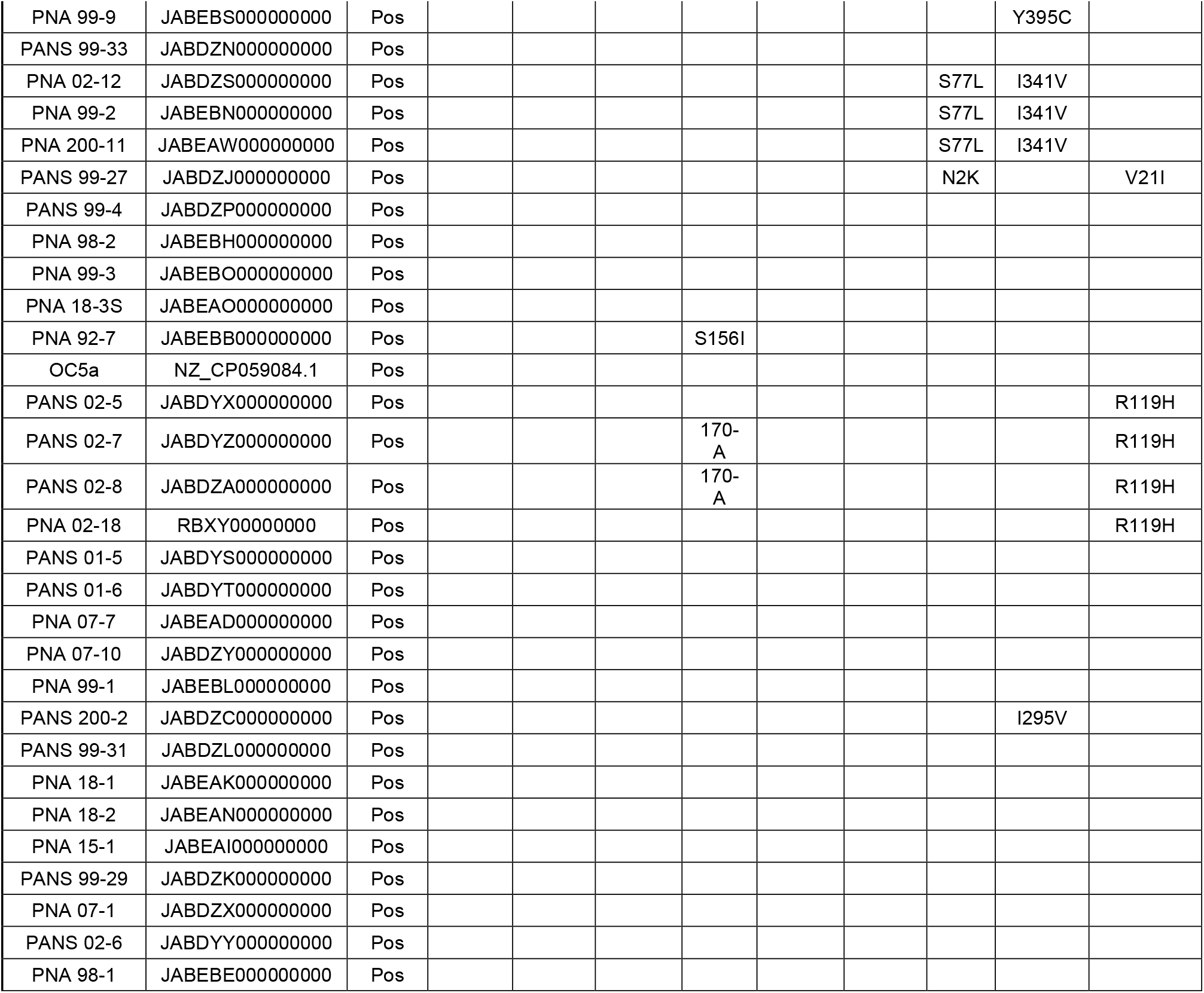
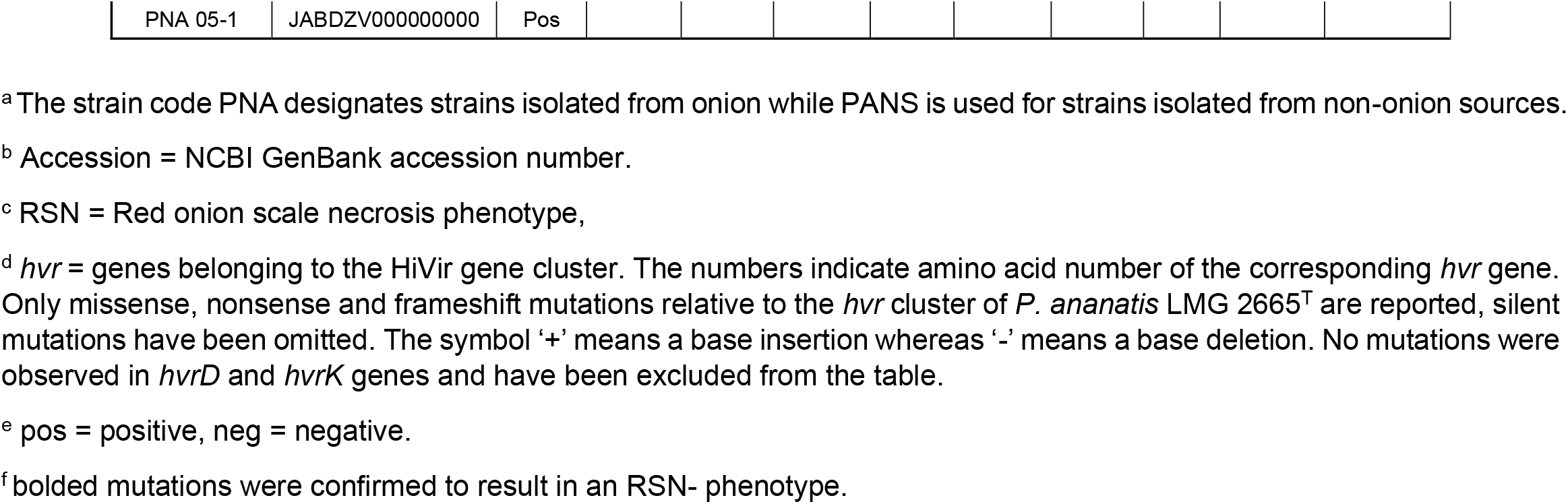
A list of mutations found in the HiVir gene cluster relative to the RSN+ Type strain *P. ananatis* LMG 2665^T^.

### The *hvrA* L216S and I260K missense mutations result in lack of RSN

To experimentally test whether D154G, L216S, I260K missense mutations found in the *hvrA* gene negatively impact HiVir-mediated RSN, the copies of mutant *hvrA* gene (*hvrA*_D154G_, *hvrA*_L216S_, *hvrA*_I260K_) was exchanged with *hvrA* gene of the RSN+ WT *P. ananatis* 97-1R. Upon acquiring *hvrA*_L216S_ and *hvrA*_I260K_, previously RSN+ *P. ananatis* 97-1R could no longer cause RSN, but no RSN phenotype change was observed in the *hvrA*_D154G_ carrying 97-1R strain (Figure 3A). The amino acid leucine (216) and isoleucine (260) were found in the conserved regions of phosphoenolpyruvate mutase of both *P. ananatis* and a phylogenetically distant species *Photorhabdus laumondii* strain TT01; however, no conservation of aspartate (154) was observed (Figure 3B). The L216S is a common mutation found in RSN-deviant *P. ananatis* strains PANS 02-1, PNA 07-14 and PNA 98-11. By replacing *hvrA*_L216S_ with *hvrA*_WT_, all three strains regained positive RSN phenotype (Figure 3C).

**Figure 3.**
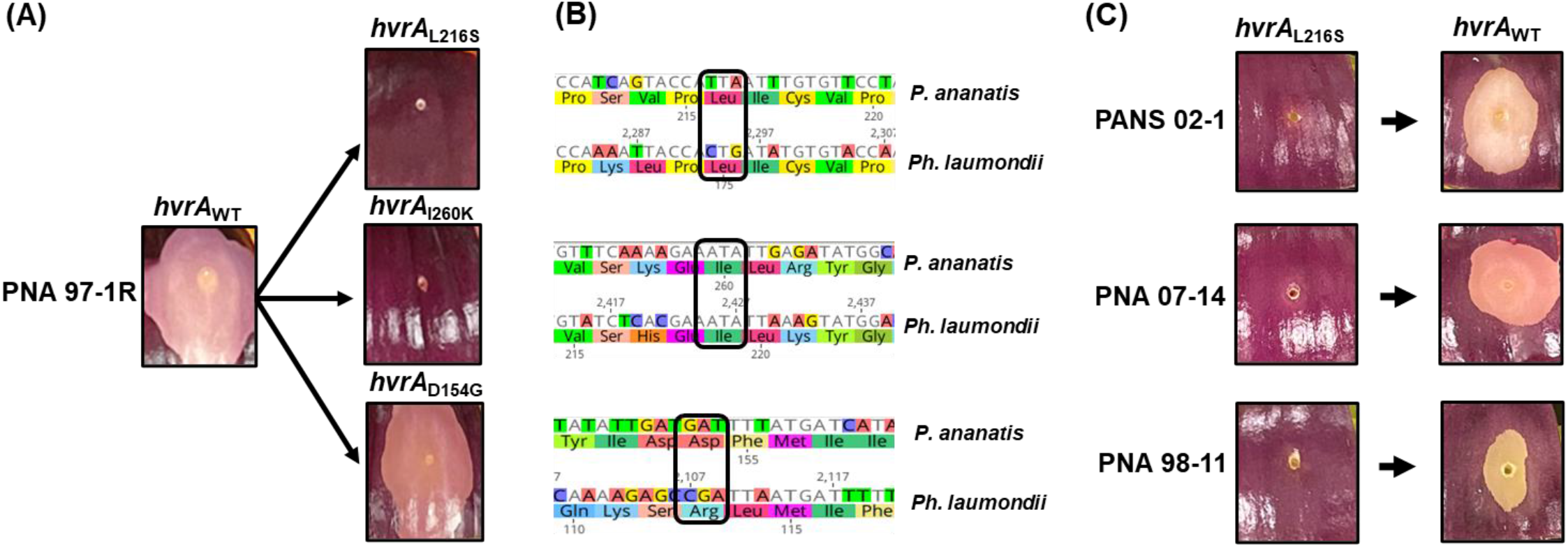
**(A)** The single nucleotide polymorphisms found in *hvrA* gene leads to leucine to serine change at amino acid 216 (L216S), isoleucine to lysine at amino acid 260 (I260K) and aspartate to glycine at amino acid 154 (D154G). The *hvrA*_L216S_ and *hvrA*_I260K_ replacement of the *hvrA* (*hvrA*_WT_) in wildtype *P. ananatis* PNA 97-1 results in the abolishment of red onion scale necrosis (RSN) whereas no change was observed in wildtype strain with *hvrA*_D154G_. **(B)** Conservation of amino acids 216 and 260 was seen when translated sequence of *Pantoea ananatis* PNA 97-1 *hvrA*/*pepM* gene was compared to that of a phylogenetically distant *Photorhabdus laumondii* subsp. *laumondii* TT01 whereas amino acid 154 was variable. **(C)** Replacing *hvrA*_L216S_ with *hvrA*_WT_ restores RSN in the phenotypically deviant *P. ananatis* strains such as PANS 02-1, PNA 07-14 and PNA 98-11.

### Double complementation of *hvrA* and *hvrC* rescues RSN in deviant *P. ananatis* strains

In addition to *hvrA*_L216S_ mutation, deviant strains PANS 04-2 and PNA 07-13 also have distinct frameshift mutations caused by the insertion of extra adenine and thiamine (*hvrC*_3+A_ and *hvrC*_32+T_, respectively) in *hvrC* gene. The co-transformation of two independent plasmids expressing *hvrA*_WT_ and *hvrC*_WT_ into the strains PANS 04-2 and PNA 07-13 rescued their RSN phenotypes (Figure 4). Furthermore, to ensure that *hvrB*_V98L_ does not contribute to the loss of RSN in PANS 04-2, *hvrB* of the WT *P. ananatis* was exchanged with *hvrB*_V98L_. Despite this exchange, WT *P. ananatis* carrying *hvrB*_V98L_ still caused RSN (Figure 4A), suggesting that V98L change in *hvrB* gene was not a RSN inactivating mutation.

**Figure 4.**
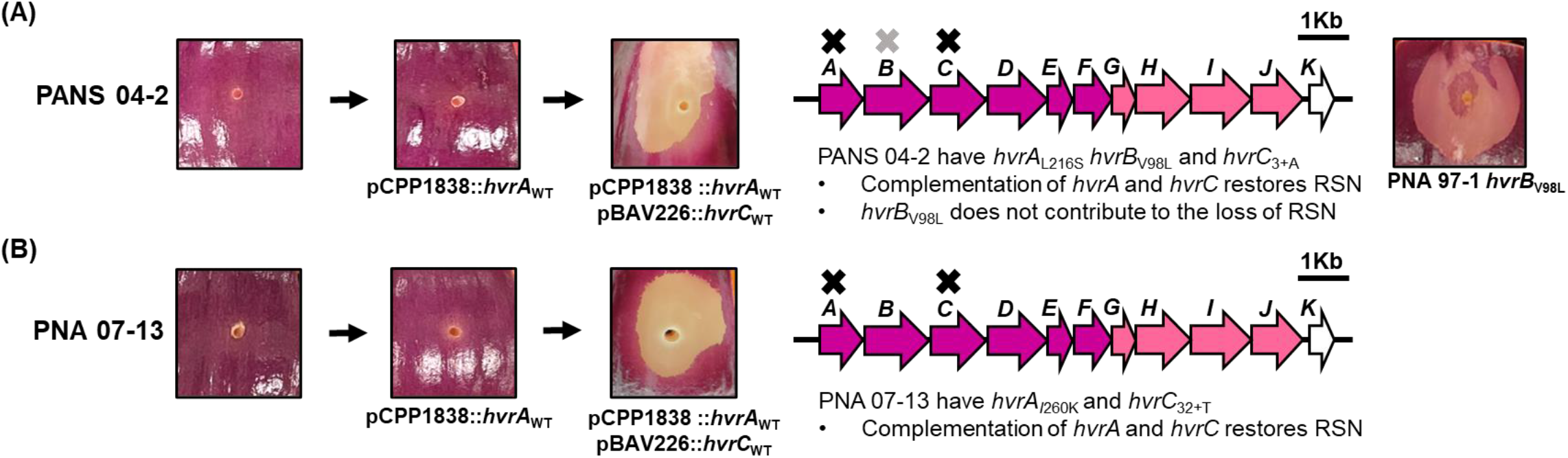
Double complementation restores red onion scale necrosis (RSN) in deviant *Pantoea ananatis* strains PANS 04-2 and PNA 07-13. **(A)** PANS 04-2 possesses three mutations (*hvrA*_L216S_, *hvrB*_V98L_, *hvrC*_3+A_) of which *hvrA*_L216S_ and *hvrC*_3+A_ are HiVir-mediated RSN inactivating mutations. Upon trans-complementation of both *hvrA* and *hvrC* genes, PANS 04-2 was able to cause RSN. The replacement of *hvrB*_WT_ with *hvrB*_V98L_ in the wildtype *P. ananatis* PNA 97-1R background shows that valine to leucine change at amino acid 98 of the *hvrB* gene does not contribute to the loss of RSN in PANS 04-2. **(B)** Similarly, PNA 07-13 have two RSN-inactivating mutations *hvrA*_I260K_ and *hvrC*_32+T_ as single complementation of *hvrA* gene with pCPP1838::*hvrA*_WT_ was not sufficient to rescue RSN phenotype. However, double complementation of *hvrA* and *hvrC* in PNA 07-13 restored RSN. The arrows represent *hvr* genes on HiVir cluster. The magenta arrows indicate essential *hvr* genes in HiVir-dependent RSN, pink = partially contributing and white = non-essential. Black cross (**X**) represents RSN-inactivating mutation whereas grey cross (**X**) indicates non RSN-inactivating mutation.

### The *hvrF* E70* mutation is a RSN inactivating mutation in *P. ananatis* PANS 99-32

The HiVir gene cluster of the deviant PANS 99-32 strain contains three SNPs (Table 1, Figure 5). As the *hvrA*_D154G_ mutation was established as an insufficient mutation to abolish HiVir-mediated RSN (Figure 3A), the non-sense mutation of *hvrF*_E70*_ and V116I change in *hvrI* were further investigated. Transformation of PANS 99-32 with pBS46::*hvrF*_WT_ allowed the strain to form a small necrotic lesion on onion scale (Figure 5A), indicating a partial complementation of *hvrF*_E70*_ with *hvrF*_WT_. A full complementation of RSN phenotype was achieved by the deletion of chromosomal copy of mutant *hvrF*_E70*_ and trans-complementation of *hvrF*_E70*_ with *hvrF*_WT_ (Figure 5B). This phenotypic restoration also suggests that *hvrI*_V116I_ does not contribute to the loss of RSN.

**Figure 5.**
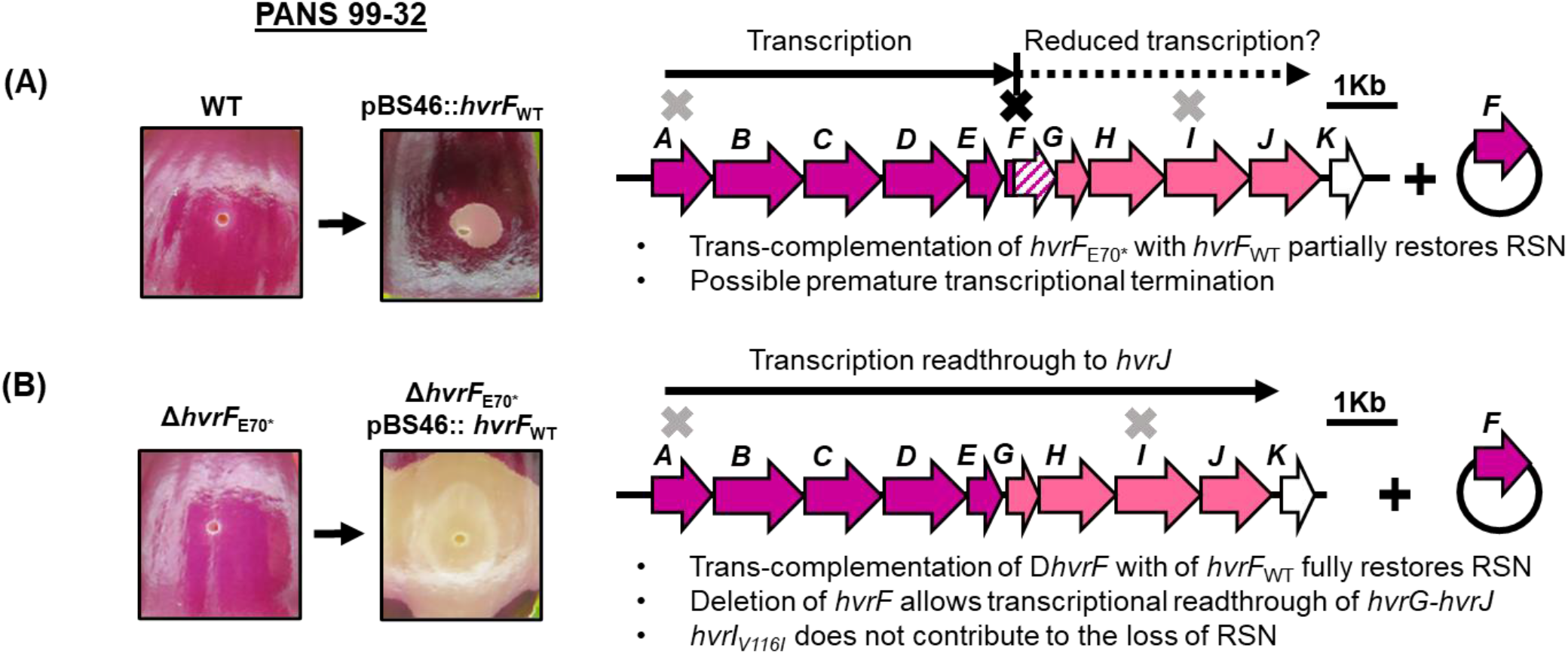
A deviant *Pantoea ananatis* strain PANS 99-32 have *hvrA* _D154G_, *hvrF*_E70*_ and *hvrI*_V116I_ mutations of which *hvrA*_D154G_ was found to be neutral as indicated by the grey cross (**X**). **(A)** Providing PANS 99-32 with a functional copy of *hvrF*_WT_ on a complementing plasmid pBS46, a partial complementation of red onion scale necrosis (RSN) was observed. The partial restoration of RSN in PANS 99-32 could be caused by the polar effect of *hvrF*_E70*_ on the downstream genes via premature transcription termination. **(B)** Deletion of chromosomal copy of *hvrF*_E70*_ and trans-complementation of *hvrF* _E70*_ with *hvrF*_WT_ on the plasmid pBS46 fully rescued the RSN ability of PANS 99-32. The removal of *hvrF*_E70*_ likely allows transcriptional readthrough from *hvrG* to *hvrJ* genes that have been found to partially contribute to the RSN ability of HiVir carrying *P. ananatis* (pink arrows). The RSN rescue by deletion and trans-complementation of *hvrF*_E70*_ also shows that *hvrI*_V116I_ does not contribute to the inactivation of RSN in PANS 99-32, indicated by the grey cross (**X**). The black cross (**X**) represents RSN-inactivating mutation.

### Cell-free spent medium from a P_tac_-driven HiVir strain can cause RSN and cell-death in *Nicotiana tabacum*

To purify and characterize pantaphos, Polidore et al. (2021) generated a *P. ananatis* strain in which the HiVir genes were driven from a heterologous P_tac_ promoter introduced into the chromosome by single cross-over (Figure 6A). We mimicked this genetic approach to produce cell-free spent supernatant from HiVir-induced strains to test for killing activity on onion scales and tobacco leaves. We recovered cell-free spent media of WT *P. ananatis* PNA 97-1R, HiVir-inducible (PNA 97-1R-SNARE::P_tac_*hvr*) and *hvrF* deletion mutant (PNA 97-1R Δ*hvrF*-SNARE::P_tac_*hvr*) as a negative control (Figure 6A). Spent media were applied to the onion scale punctures as done previously. The IPTG-induced spent medium of HiVir-inducible strain caused a *P. ananatis*-characteristic necrosis of the red onion scales at 4 dpi whereas no symptoms were observed in the scales inoculated with the spent media of the WT or HiVir-inducible, *hvrF* deletion mutant negative control strain (Figure 6B). In the absence of induction, the PNA 97-1R, PNA 97-1R-SNARE::P_tac_*hvr* spent medium also failed to produce an RSN phenotype (Figure 6A under ‘Non-IPTG induced’).

**Figure 6.**
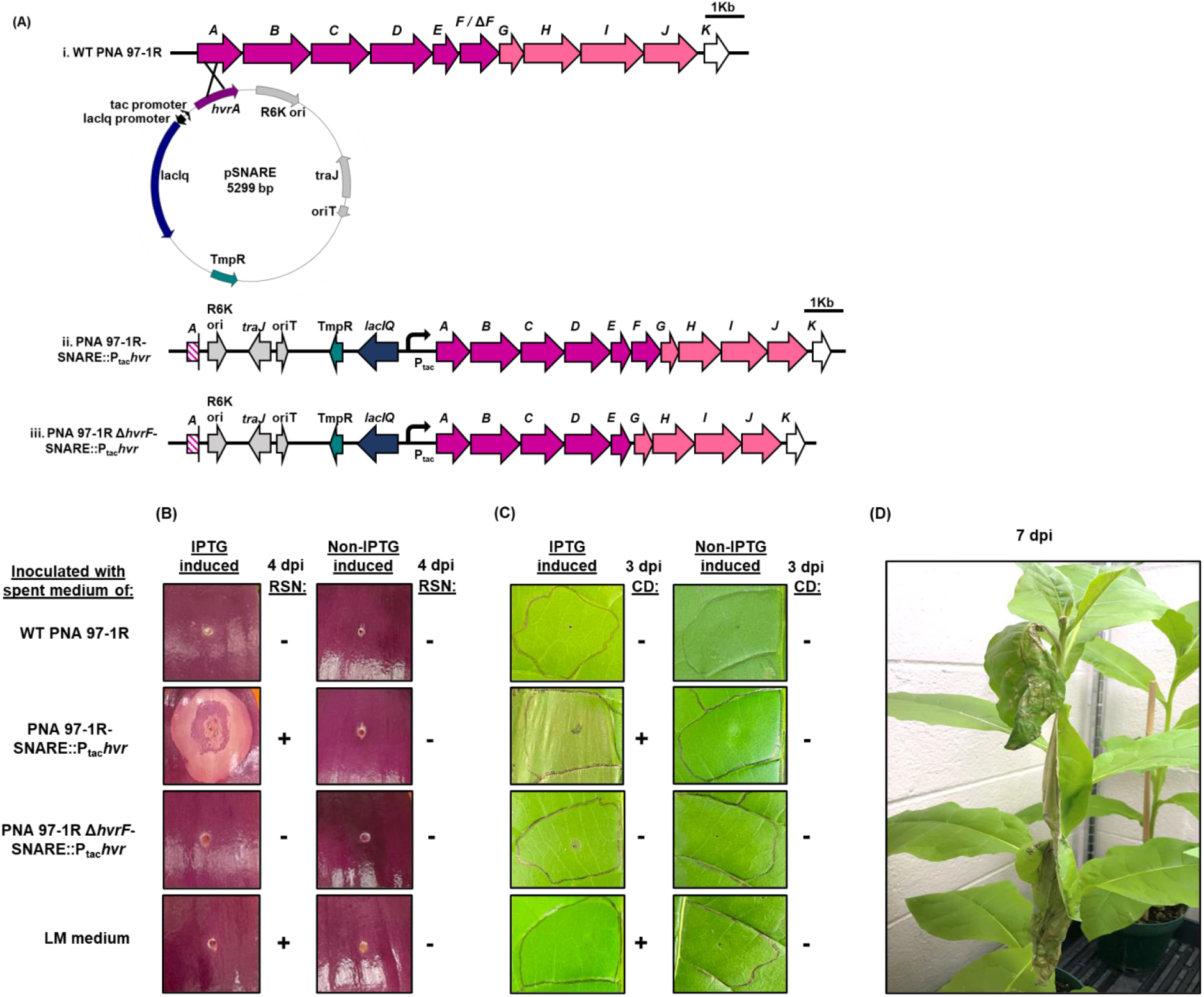
**(A)** Construction of IPTG-inducible HiVir strain of *Pantoea ananatis* PNA 97-1R. **i:** The plasmid pSNARE carrying partial *hvrA* gene (450 bp) downstream of the IPTG-inducible promoter (P_tac_) was introduced into the wildtype (WT) and *hvrF* deletion mutant (Δ*hvrF*) strains of *P. ananatis* PNA 97-1R via conjugation with *Escherichia coli* RHO5 (donor). The pSNARE is integrated into the chromosome of PNA 97-1R strains by homologous recombination at *hvrA* gene, resulting in IPTG-inducible HiVir gene cluster in both **ii**. WT (PNA 97-1R-SNARE::P_tac_*hvr*) and **iii**. *hvrF* deletion mutation (PNA 97-1R Δ*hvrF*-SNARE::P_tac_*hvr*) background. **(B)** The red onion scale necrosis (RSN) and **(C)** Tobacco (*Nicotiana tabacum*) infiltration assays using the spent media of IPTG-induced and non-induced cell-free spent media of wildtype (WT) *Pantoea ananatis* PNA 97-1R, PNA 97-1R-SNARE::P_tac_*hvr* and Δ*hvrF* PNA 97-1R-SNARE::P_tac_*hvr* were **(A)** drop-inoculated (10 µl) on red onion scales and **(B)** syringe infiltrated (100 µl) into tobacco leaves. Sterile LM broth was used as a medium control and images were taken at 4 days post inoculation (dpi) and 3 dpi, respectively. Positive RSN and cell death (CD) symptoms were observed in onion scales and tobacco leaves infiltrated with IPTG-induced spent medium of PNA 97-1R-SNARE::P_tac_*hvr*. **(C)** Tobacco plant showing extended leaf spot symptoms in upper, non-infiltrated leaves at 7 dpi.

When induced spent medium of the HiVir-inducible strain was infiltrated into tobacco leaf panels, cell death developed in the infiltrated area at 3 dpi (Figure 6C). However, cell death was not produced by the spent media of WT nor HiVir-inducible Δ*hvrF* deletion mutant strains, irrespective of induction. Surprisingly, over the course of seven days, the infiltrated tobacco plant developed necrotic spots outside of the infiltrated leaf panel and then on leaves that were not infiltrated with the spent media (Figure 6D). The symptoms progressed upwards to leaves directly above the infiltrated leaf towards the apex of the plant, eventually leading to the development of malformed leaves and the formation of a lateral stem.

### Application of HiVir-induced spent medium is sufficient to restore *in planta* bacterial populations of *hvr* gene mutant strains

The HiVir-induced toxin <u>s</u>pent <u>m</u>edium (**tsm**) of HiVir-inducible (PNA 97-1R-SNARE::P_tac_*hvr*) and *hvrF* <u>m</u>utant <u>s</u>pent <u>m</u>edium (**msm**) of HiVir-inducible Δ*hvrF* deletion mutant (Δ*hvrF*-SNARE::P_tac_*hvr*) *P. ananatis* strains were used in the co-inoculation of WT PNA 97-1R, Δ*hvrA*, Δ*hvrD* and *hvrA*_L216S_ strains. The RSN symptoms were observed in the scales inoculated with WT and co-inoculated with tsm whereas scales infected with RSN-mutant Δ*hvrA*, Δ*hvrD* and *hvrA*_L216S_ strains and co-inoculated with msm did not show any symptoms (Figure 7A). Consistent with the absence of RSN, *in planta* bacterial loads of Δ*hvrA*, Δ*hvrD* and *hvrA*_L216S_ strains with or without msm were significantly lower than that of the WT. Interestingly, *in planta* bacterial populations of the mutant strains were complemented to that of the WT-level when inoculated with tsm (Figure 7B). No bacterial growth was present in scales inoculated with just spent media and water controls.

**Figure 7.**
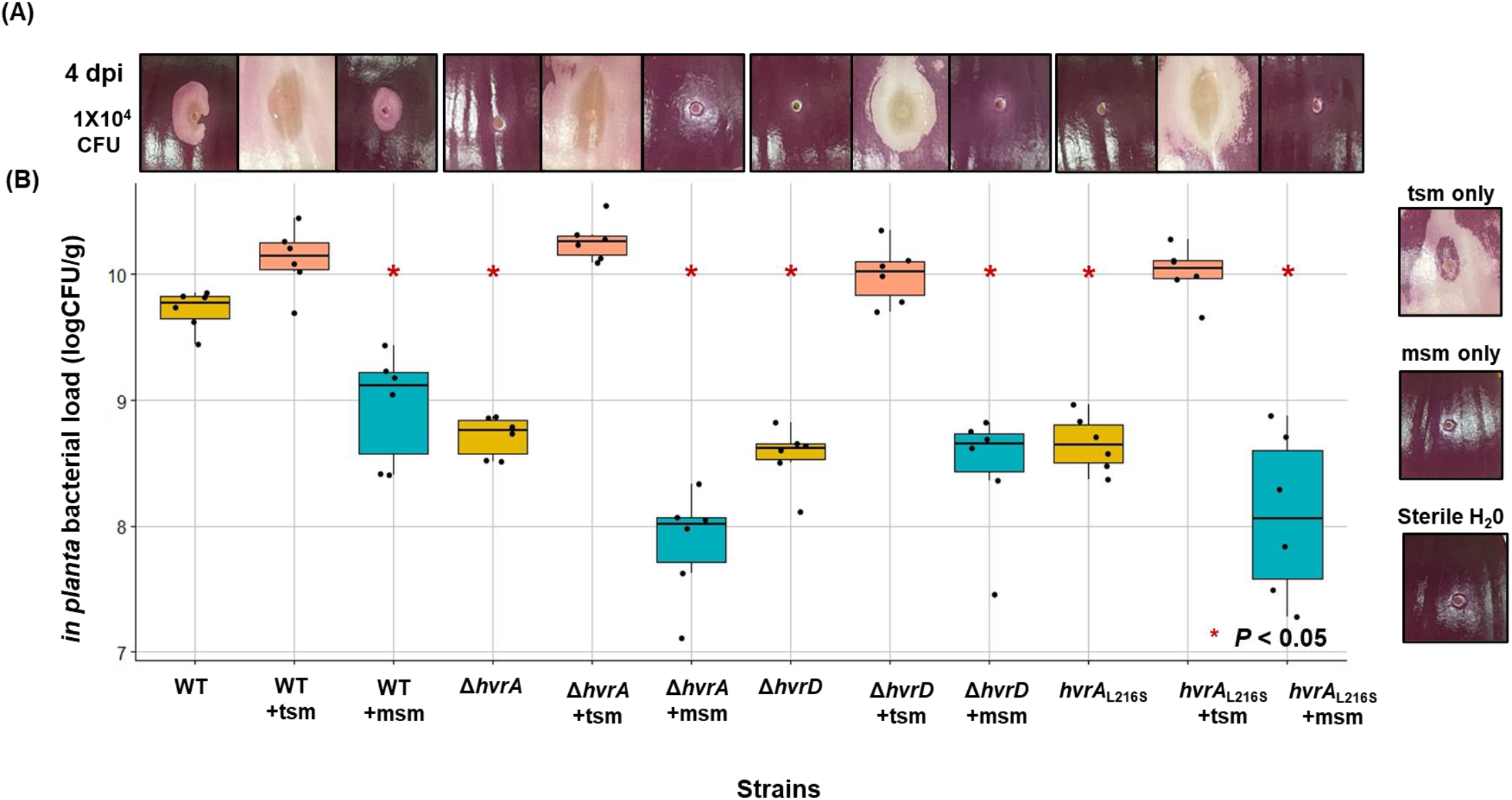
**(A)** The red onion scales inoculated with 1×10^4^ CFU of **WT** = wildtype, **Δ*hvrA*, Δ*hvrD*** and ***hvrA***_**L216S**_ mutation carrying *Pantoea ananatis* PNA 97-1R strains. The strains were simultaneously inoculated with 10 µl of **tsm** = HiVir-induced toxin <u>s</u>pent <u>m</u>edium of HiVir-inducible *P. ananatis* (PNA 97-1R-SNARE::P_tac_*hvr*) or **msm** = *hvrF* <u>m</u>utant <u>s</u>pent <u>m</u>edium (**msm**) of HiVir-inducible Δ*hvrF* deletion mutant *P. ananatis* (PNA 97-1R Δ*hvrF*-SNARE::P_tac_*hvr*). The scale images were taken at 4 days post inoculation (dpi). **(B)** The *in planta* bacterial load of each inoculum at 4 dpi is compared to that of the WT and the significant (*P* < 0.05) reduction in population size is indicated by the asterisk (*) in a box plot lot. No bacterial load was present in the scales inoculated with cell-free tsm, msm and sterile water. This experiment was repeated twice.

## Discussion

Through the creation of in-frame *hvr* gene deletion mutants, the genetic requirement of individual *hvr* genes in HiVir-mediated onion RSN and foliar necrosis was investigated. In addition to *hvrA* (Asselin et al. 2018; Stice et al. 2020), we determined that the genes, *hvrB* to *hvrF*, make an essential contribution to the HiVir-mediated RSN and foliar symptoms whereas *hvrG* to *hvrJ* genes make partial contributions. We did not observe any contribution of the *hvrK* gene to any onion necrosis associated phenotype. In both RSN and foliar assays, essential gene deletion mutants (Δ*hvrB*, Δ*hvrC*, Δ*hvrD*, Δ*hvrE*, Δ*hvrF*) failed to cause foliar lesions. The absence of RSN correlated with the significant reduction *in planta* of populations of the Δ*hvrB*, Δ*hvrC*, Δ*hvrD*, Δ*hvrE* and Δ*hvrF* strains in red onion scales (Figure 1). Except for the *hvrF* gene, the other four essential genes had been predicted to encode enzymes that are directly involved in the synthesis of pantaphos. The essential functions of these enzymes are likely to be the reason for their conservation across the different types of HiVir-like biosynthetic gene cluster (Polidore et al. 2021). On the other hand, the role of *hvrF* in the biosynthesis of pantaphos is currently unclear. Although the protein product of *hvrF* gene (O-methyltransferase) is not part of the proposed pantaphos pathway, the deletion of *hvrF* resulted in the loss of pathogenicity in *P. ananatis* as well as the loss of phytotoxicity of the spent medium produced by the HiVir-inducible Δ*hvrF P. ananatis* strain. The *hvrF*-encoded O-methyltransferase has been predicted to be involved in the resistance against the undesired trans-isomer of the side product that may form during pantaphos biosynthesis, which could be critical for its biosynthesis (Polidore et al. 2021).

The onion scales inoculated with Δ*hvrI* at 1×10^4^ CFU developed scale lesion that appear reduced compared to the lesion produced by wildtype whereas weak or no symptoms were seen in the scales inoculated with Δ*hvrG*, Δ*hvrH* and Δ*hvrJ* at the same concentration. Interestingly, the RSN phenotype could be rescued by inoculating scales with the increased concentrations (1×10^10^ CFU) of Δ*hvrG* (N-acetyltransferase), Δ*hvrH* (ATP-grasp protein) and Δ*hvrJ* (unknown). It has been proposed that the protein products of *hvrG* and *hvrH* may catalyze peptidic derivative of pantaphos for self-resistance against pantaphos or pantaphos intermediates (Polidore et al. 2021). Polidore et al. (2021) detected three phosphonic compounds of which compound 1 was designated as pantaphos, compound 2 as a precursor of pantaphos (2-phosphonomethylmaleate) in the proposed pantaphos biosynthesis pathway and compound 3 that is yet to be characterized. The presence of RSN in Δ*hvrI* (MFS transporter protein) inoculated onion scales could be explained by the availability of alternative transporters or the release of toxins from lysed bacteria. It is possible that with increased *in planta* population size, more pantaphos could be released into onion environment despite the absence of *hvrI*-encoded transporters. Similarly, although reduced, the foliar lesions caused by the Δ*hvrG*, Δ*hvrH* and Δ*hvrI* indicate production and release of pantaphos by the mutant cells. It will be interesting to determine whether increased inoculum concentration of Δ*hvrJ* will result in the foliar lesion as it has been the case for the RSN assay. Lastly, the deletion of *hvrK* (flavin reductase) did not affect the pathogenicity of *P. ananatis* nor *in planta* bacterial population. The activity of HvrK in HiVir-mediated RSN is possibly dispensable and proposed conversion of 2-phosphonomethylmaleate to pantaphos (Polidore et al. 2021) may be completed by an enzyme other than HvrK.

Pantaphos produced by HiVir gene cluster is likely to be a broad-spectrum toxin. Presently, the target of phosphonate toxin in plant is unknown yet the bioactivity of this phosphonate compound has been shown to be effective in both monocot plant: onion (Asselin et al. 2018) and eudicot Rosid: mustard seedling (Polidore et al. 2021) and Asterid: tobacco plants. The phytotoxicity of pantaphos against the mustard seedlings was comparable to that of the glyphosate, a phosphonate based systemic herbicide (Polidore et al. 2021). Glyphosate is a broad-spectrum toxin that inhibits the activity of a key plant enzyme 5-enolpyruvylshikimate-3-phosphate synthase (EPSP) by acting as a structural analog of an EPSP substrate (Schӧnbrunn et al. 2001). As a structural analog, pantaphos might inhibit the activity of an important enzyme that is part of a pathway that is likely conserved in different plant clades. Furthermore, development of cell death symptoms in tobacco plants in an ascending manner by HiVir-induced toxin-containing spent medium (Figure 6D) indicates potential systemic movement of the toxin unlike the confined cell death symptom observed in tobacco leaf panels caused by the infiltration of primed *P. ananatis* (Carr et al. 2010; Kido et al. 2010; Asselin et al. 2018). Identification of the pantaphos target could assist in efforts to breed onions that would be resistant *P. ananatis*.

Phenotypically deviant strains of *P. ananatis*, genomically HiVir+ yet phenotypically RSN-, had been previously noted by Agarwal et al., Polidore et al. and Stice et al. (2021). In six of these strains, we detected mutation in RSN essential genes of HiVir, especially in the *hvrA* gene, which encodes phosphoenolpyruvate mutase (PepM) that catalyzes the first chemical reaction in the proposed pantaphos biosynthesis pathway (Polidore et al. 2021). The missense mutations such as L216S and I260K on conserved or structurally important amino acids could potentially be detrimental to protein folding and catalytic activity of PepM. Moreover, inactivation of this first enzyme would halt the entire predicted pantaphos biosynthesis pathway possibly resulting in the lack of pataphos production which is consistent with the previous (Asselin et al. 2018, Stice et al. 2020) and current findings where deletion or inactivation of *hvrA* in *P. ananatis* failed to cause RSN symptoms unless complemented with the wildtype copy of *hvrA* gene.

Other RSN-inactivating mutations were frameshift and nonsense mutations, which resulted in the coding of premature stop codons in *hvrC* gene of PANS 04-2 and PNA 07-13 and *hvrF* in PANS 99-32, respectively. As PANS 04-2 and PNA 07-13 carried distinct RSN-inactivating mutations in both *hvrA* and *hvrC* genes, double complementation of these genes was necessary for the full restoration of RSN (Figure 4). However, trans-complementation of one RSN-inactivating mutation *hvrF*_E70*_ with pBS46::*hvrF*_WT_ in PANS99-32 resulted in partial phenotypic restoration of onion necrosis (Figure 5A). Only when chromosomal *hvrF*_E70*_ was removed, trans-complementation with pBS46::*hvrF*_WT_ in PANS99-32 resulted in full RSN rescue (Figure 5B). This observation is consistent with the polar effect of *hvrF*_E70*_ on downstream *hvrG* to *hvrJ* genes. It is possible that premature termination of translated amino acid at *hvrF*_E70*_ exposes mRNA naked, allowing transcription terminator factor such as rho to bind and cause early transcription termination of downstream *hvr* genes. However, deletion of this ‘signal’ could allow transcriptional readthrough of *hvrG* to *hvrJ* genes that still partially contribute to the onion necrosis phenotype.

The co-inoculation experiment of HiVir-induced toxin-containing <u>s</u>pent <u>m</u>edium (**tsm**) or *hvrF* <u>m</u>utant <u>s</u>pent <u>m</u>edium (**msm**) with the RSN negative Δ*hvrA*, Δ*hvrD* and L216S mutant *P. ananatis* PNA 97-1R strains revealed that the exogenous application of the toxin or tsm was sufficient for the establishment of WT-level *in planta* bacterial populations in HiVir-inactivated strains (Figure 7). As previously suggested by Stice et al. (2020), *P. ananatis* displays a necrotrophic lifestyle that is mediated by HiVir and is further facilitated by the plasmid-borne *alt* (<u>al</u>licin tolerance) gene cluster, which confers tolerance to thiosulfinates released by necrotized onion tissue (Stice et al. 2020). Detoxification of organosulfur compounds is crucial for the proliferation of *P. ananatis* especially in onion bulb environment. In fact, HiVir+ *alt*-*P. ananatis* strains had a significantly lower incidence of bulb rot than HiVir+ *alt*+ strains when bacterial progression was assessed from onion neck to bulb tissue (Stice et al. 2021). It is interesting to note that deviant *P. ananatis* strains lack *alt* (Agarwal et al. 2021) and it is likely that they would be at a selective disadvantage when confronted with necrotic onion environments. Therefore, inactivation of HiVir by accumulating deleterious mutations in essential *hvr* genes would prevent these strains from producing toxic secondary metabolite such as pantaphos and allow them to adopt a non-pathogenic or epiphytic lifestyle.

The HiVir-*alt* strategy may be common a virulence strategy shared among species of *Pantoea* implicated in Onion Center Rot disease. *P. allii* (Edens et al. 2006; Brady et al. 2011), and *P. agglomerans* (Hattingh and Walters, 1981; Edens et al. 2006; Tho et al. 2015) are often associated with Onion Center Rot and these members of *Pantoea* share an *P. ananatis*-like HiVir cluster (Figure S1) and an *alt* cluster (Stice et al. 2021). The role of HiVir cluster in onion pathogenicity of *P. allii* and *P. agglomerans* and the regulation of the HiVir cluster in the species of *Pantoea* remains to be addressed. Furthermore, identification of the molecular target of the phosphonate toxin in onion plant will be of valuable information for the development of toxin-resistant onion.

## Acknowledgement

We would like to acknowledge Dr. J. Asselin and Dr. B. Vinatzer for graciously providing plasmids pCPP1383::*hvrA* and pBAV226, respectively, and Dr. P. Stodghill and Dr. J. Asselin for providing helpful comments and critiques regarding the preparation of this manuscript.

## Funding

This work is supported by Specialty Crops Research Initiative Award 2019-51181-30013 from the USDA, National Institute of Food and Agriculture to BD and BK. Any opinions, findings, conclusion, or recommendations expressed in this publication are those of the author(s) and do not necessarily reflect the view of the U.S. Department of Agriculture.

## Notes

### Competing Interest Statement

The authors have declared no competing interest.

