## Supplementary Figure 1 for "The genetic requirements for HiVir-mediated onion necrosis by *Pantoea ananatis*, a necrotrophic plant pathogen"

(A)

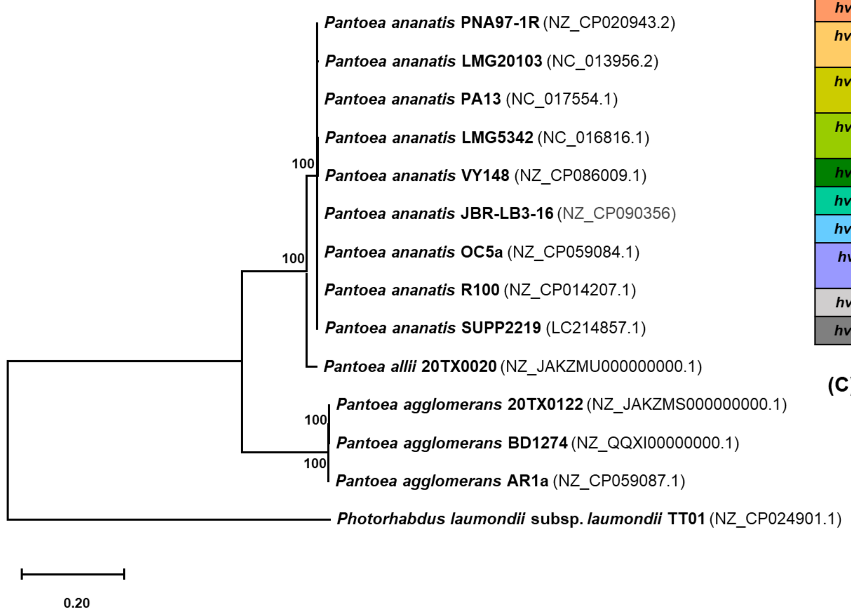


**(B) Putative functions of HiVir and HiVir-like genes in *Pantoea* and *Photorhabdus laumondii***

| *Pantoea* | | *Photorhabdus* | |
|---|---|---|---|
| Genes | Annotations | Genes | Annotations |
| *hvrA* | Phosphoenolpyruvate mutase | plu1865 | Phosphoenolpyruvate mutase |
| *hvrB* | Flavin-dependent monooxygenase | plu1866 | Nitrilotriacetate monooxygenase |
| *hvrC* | Phosphomethylmalate synthase | plu1867<br>plu1868 | Pseudogene:<br>Homocitrate synthase |
| *hvrD* | Isopropylmalate dehydratase-like protein large subunit | plu1869 | 3-isopropylmalate dehydratase, large subunit |
| *hvrE* | Isopropylmalate dehydratase-like protein small subunit | plu1870 | 3-isopropylmalate dehydratase, small subunit |
| *hvrF* | O-methyltransferase | plu1871 | Methyltransferase |
| *hvrG* | GNAT family acetyltransferase | plu1872 | Acetyletransferase |
| *hvrH* | ATP-grasp protein | plu1873 | Similarities with NikS protein |
| *hvrI* | MFS-transporter protein | plu1874 | Similarities with Nikkomycin biosynthesis carboxylase |
| *hvrJ* | Unknown | plu1875 | Integral membrane efflux protein |
| *hvrK* | Flavin reductase | plu1876 | Unknown |

(C)

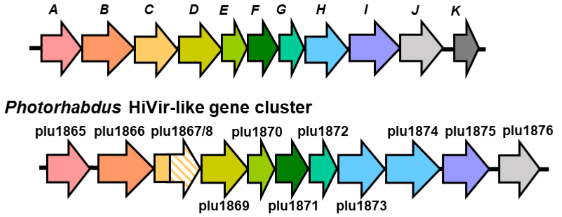


**Supplementary figure 1. (A)** A maximum likelihood tree based on HiVir and HiVir-like gene cluster retrieved from public genomes of *Pantoea* species and *Photorhabdus laumondii* strain TT01. The genome accession numbers are indicated in the brackets next to the species name. The general time reversible (GTR) nucleotide substitution was selected with invariable sites (I) of 0.195 and gamma distribution (G) of 3.054. A bootstrap value (%) after 1000 replicates are shown at the nodes and *Photorhabdus laumondii* subsp. *laumondii* strain TT01 was used as an outgroup. A scale bar shows a nucleotide substitution rate per site. **(B)** A table showing predicted annotations of *hvr* genes and HiVir-like genes of *Photorhabdus laumondii*. The coloring is based on the annotation. **(C)** The organization and annotations of HiVir-like gene cluster found in *Photorhabdus laumondii* closely resembles that of *Pantoea* species with a few exceptions indicated by white arrows. A putative *hvrK* encoded flavin reductase is absent in *Photorhabdus laumondii* cluster. The gene clusters are not drawn to scale.
