## Supplementary Tables for "The genetic requirements for HiVir-mediated onion necrosis by *Pantoea ananatis*, a necrotrophic plant pathogen"

**Table S1**. Bacterial strains and plasmids used in cloning and mutagenesis

| Name | Description | Reference |
|---|---|---|
| **1. *P. ananatis*** | | |
| PNA 97-1R | HiVir+ RSN+, spontaneous rifampicin mutant ($Rp^R$) of *P. ananatis* PNA 97-1 | Stice et al. 2018 |
| PNA 97-1R pBS46::EV | HiVir+ RSN+, spontaneous rifampicin mutant ($Rp^R$) of *P. ananatis* PNA 97-1 harboring empty vector pBS46 ($Gm^R$) | This study |
| PNA 97-1R Δ*hvrB* | HiVir+ RSN-, a *hvrB* deletion mutant of *P. ananatis* PNA 97-1R | This study |
| PNA 97-1R Δ*hvrC* | HiVir+ RSN-, a *hvrC* deletion mutant of *P. ananatis* PNA 97-1R | This study |
| PNA 97-1R Δ*hvrD* | HiVir+ RSN-, a *hvrD* deletion mutant of *P. ananatis* PNA 97-1R | This study |
| PNA 97-1R Δ*hvrE* | HiVir+ RSN-, a *hvrE* deletion mutant of *P. ananatis* PNA 97-1R | This study |
| PNA 97-1R Δ*hvrF* | HiVir+ RSN-, a *hvrF* deletion mutant of *P. ananatis* PNA 97-1R | This study |
| PNA 97-1R Δ*hvrG* | HiVir+ RSN+, a *hvrG* deletion mutant of *P. ananatis* PNA 97-1R | This study |
| PNA 97-1R Δ*hvrH* | HiVir+ RSN+, a *hvrH* deletion mutant of *P. ananatis* PNA 97-1R | This study |
| PNA 97-1R Δ*hvrI* | HiVir+ RSN+, a *hvrI* deletion mutant of *P. ananatis* PNA 97-1R | This study |
| PNA 97-1R Δ*hvrJ* | HiVir+ RSN+, a *hvrJ* deletion mutant of *P. ananatis* PNA 97-1R | This study |
| PNA 97-1R Δ*hvrK* | HiVir+ RSN+, a *hvrK* deletion mutant of *P. ananatis* PNA 97-1R | This study |
| PNA 97-1R Δ*hvrB* pBS46::*hvrB* | HiVir+ RSN-, a *hvrB* deletion mutant of *P. ananatis* PNA 97-1R harboring *hvrB* complementation construct, $Gm^R$ | This study |
| PNA 97-1R Δ*hvrC* pBS46::*hvrC* | HiVir+ RSN-, a *hvrC* deletion mutant of *P. ananatis* PNA 97-1R harboring *hvrC* complementation construct, $Gm^R$ | This study |
| PNA 97-1R Δ*hvrD* pBS46::*hvrD* | HiVir+ RSN-, a *hvrD* deletion mutant of *P. ananatis* PNA 97-1R harboring *hvrD* complementation construct, $Gm^R$ | This study |

| | | |
|---|---|---|
| PNA 97-1R Δ*hvrE* pBS46::*hvrE* | HiVir+ RSN-, a *hvrE* deletion mutant of *P. ananatis* PNA 97-1R harboring *hvrE* complementation construct, $Gm^R$ | This study |
| PNA 97-1R Δ*hvrF* pBS46::*hvrF* | HiVir+ RSN-, a *hvrF* deletion mutant of *P. ananatis* PNA 97-1R harboring *hvrF* complementation construct, $Gm^R$ | This study |
| PNA 97-1R Δ*hvrG* pBS46::*hvrG* | HiVir+ RSN+, a *hvrG* deletion mutant of *P. ananatis* PNA 97-1R harboring *hvrG* complementation construct, $Gm^R$ | This study |
| PNA 97-1R Δ*hvrH* pBS46::*hvrH* | HiVir+ RSN+, a *hvrH* deletion mutant of *P. ananatis* PNA 97-1R harboring *hvrH* complementation construct, $Gm^R$ | This study |
| PNA 97-1R Δ*hvrI* pBS46::*hvrI* | HiVir+ RSN+, a *hvrI* deletion mutant of *P. ananatis* PNA 97-1R harboring *hvrI* complementation construct, $Gm^R$ | This study |
| PNA 97-1R Δ*hvrJ* pBS46::*hvrJ* | HiVir+ RSN+, a *hvrJ* deletion mutant of *P. ananatis* PNA 97-1R harboring *hvrJ* complementation construct, $Gm^R$ | This study |
| PNA 97-1R Δ*hvrK* pBS46::*hvrK* | HiVir+ RSN+, a *hvrK* deletion mutant of *P. ananatis* PNA 97-1R harboring *hvrK* complementation construct, $Gm^R$ | This study |
| PNA 97-1R-pSNARE | HiVir+ RSN+, spontaneous rifampicin mutant ($Rp^R$) of *P. ananatis* PNA 97-1 with inducible HiVir gene cluster (Ptac-HiVir) | This study |
| PNA 97-1R Δ*hvrF*-pSNARE | HiVir+ RSN-, a *hvrF* deletion mutant of *P. ananatis* PNA 97-1R with inducible HiVir gene cluster (pTac-HiVir) | This study |
| **2. Deviant *P. ananatis* strains** | | |
| PANS 02-1 | HiVir+ RSN- strain, isolated from onion thrip | Stice et al. 2018 |
| PANS 04-2 | HiVir+ RSN- strain, isolated from onion thrip | Stice et al. 2018 |
| PANS 99-32 | HiVir+ RSN- strain, isolated from Florida Pusley | Stice et al. 2018 |
| PNA 07-13 | HiVir+ RSN- strain, isolated from onion | Agarwal et al. 2021 |
| PNA 07-14 | HiVir+ RSN- strain, isolated from onion | Agarwal et al. 2021 |
| PNA 98-11 | HiVir+ RSN- strain, isolated from onion | Stice et al. 2018 |

| | | |
|---|---|---|
| **3. *E. coli* strains** | | |
| DH5α | General cloning strain | Liss, 1987 |
| MaH1 | DH5α derivative, attTn7 pir116 R6K replicon plasmids | Kvitko et al. 2012 |
| RHO5 | SM10 derivative, pir116, DAP-dependent conjugation strain | Kvitko et al. 2012 |
| **4. Plasmids** | | |
| pR6KT2G | Gateway-derivative of pR6KT2 with BP clonase compatible cassette, *sacB*, $Gm^R$, *gus*, $Cm^R$ | Stice et al. 2020 |
| pDONR221 | Gateway BP clonase compatible cloning vector | Invitrogen |
| pBS46 | pBBR1-MCS5 derivative, gateway LR clonase compatible expression vector, *ccdB*, $Gm^R$ | Swingle et al. 2008 |
| pBS46::EV | Empty expression vector, $Gm^R$ | Swingle et al. 2008 |
| pSC201 | A variant of the pSCrhaB2 vector with oriR6K, rhamnose inducible promoter (pRha), $Tmp^R$ | Ortega et al. 2007 |
| pSNARE | A modified pSC201 with oriR6K, IPTG inducible promoter ($P_{Tac}$) harboring partial *hvrA* gene fragment, used for homologous recombination into the chromosome of *P. ananatis* PNA 97-1R, $Tmp^R$ | This study |
| pCPP1838::OC5a_*hvrA* | hvrA complementing plasmid (*hvrA* gene and 22 bp upstream cloned from *P. ananatis* strain OC5a), RSF1010 ori, $Gm^R$, $Km^R$ | Asselin et al. 2018 |
| pBAV226 | Gateway LR clonase compatible expression vector, p15A ori, $Tet^R$, $Gm^R$ | Vinatzer et al. 2006 |

**Table S2.** A list of dsDNA synthesized for the deletion mutation and genetic engineering of *P. ananatis* PNA 97-1 HiVir gene cluster

| Name | Sequence |
|---|---|
| **delhvrB** | ACAAGTTTGTACAAAAAAGCAGGCTGTTTATATTGATGATTTTATGATCATAGCCAGACTTGAAAGTCTTATTGCAGGGTTC<br>GACGTAGAACATGCACTCGAACGTGCCGACGCATACGTCGAAGCCGGGGCAGACGGAATTATGATTCATAGTTGTAAGAA<br>GACTCCGGATGAGGTTTTCTTATTCAGTACGAAATTTCGGAAAAAATATCCATCAGTACCATTAATTTGTGTTCCTACTACTT<br>ATTCTGCAACCAGCAACAGAGAACTCAGTGAAGCGGGTTTTAACGTGATCATTTATGCAAACCATATGCTCAGGGCTGCTT<br>ATAAAGCAATGGAAAATGTTTCAAAAGAAATATTGAGATATGGCAGGACGGCAGAGATAGAAAAATCTTGCATGAGTGTAA<br>AGGAAATTATTTCACTGATTCCTTAAAGAGAATGACCATGAAAAAAAAAGAGATGATAATAGGTGCCCCCGGGAGACCTAC<br>TAATAAATTGTTTAATCAATAATGTATTGCGAGGAAATATAAAATGCTTAATAAAAATCTGATTCTTGAAGATACCACTTTACG<br>TGATGGTGAGCAGGCGCCAGGTGTTGCATTTACACCAGAGCAAAAAGTAGAAATTTTTTATCTACTTGCAAATATGGGCGT<br>TAAATGGATCGAAGCCGGAATACCTGCGATGAAGGGTGATGAAGTAAAGGCTCTGTCGGAAATGTTAGAGAGAAAAAATG<br>AAATTAACATCATCGCGTGGAACCGAGGCGTGCTTGAAGACATTGAGTACAGTATCTCACTTGGATTCAAAGCGGTGCATA<br>TCGGGCTACCGACTTCAGCTATCCATTTAGAGAAAAGCGTTAAGAAAGATAAGTCCTGGCTTGTAAAGACGGCTTCAGATT<br>TAGTTAAGTTTGCCAAAGACAAAGGGATGTTTGTTACCCAGCTTTCTTGTACAAAGTGGT |
| **delhvrC** | ACAAGTTTGTACAAAAAAGCAGGCTTTACTGGGTGGGGTTGATTTAAAAAACAATGATCCTGAAGGACCATTACCAGATTTA<br>CCGAAATCTAATGGTAATCAGAGTAGACAAAAAATTATTATCGATTTAGCTCGAAAAGAAAAACTATCTATCAAGCAACTTTA<br>TGAAAAAATAATCATTTCAAGAGGACATTATACATTTACAGGTTCCTATCAAGATTTAGCAGATGAGATGATTAAGTGGGTT<br>GAAAATGAAGCATGTGATGGTTTCAACATTATGCCTCCTCTCATGCCTGAATCTCTTATTAATCTTTTCGATCATGTCATTCC<br>ACTTATTCAGGCAAGAGGATGGTATAAAAAATCATATTCTACTGGAACATTAAGAGAAAAACTGGGGCTTAAAAGACCTACT<br>AATAAATTGTTTAATCAATAATGTATTGCGAGGAAATATAAAATGCTTAATAAAAATCTGATTCCCGGGGATCTGTATACCGC<br>GGGGTAAACAGGTATGAAAAGTAATCAGCCGATTGTTAACCAGATCATTGCGTCACACAGTGGCAGAGGTCAGGTTTCAG<br>CAGGTGAACTGATCACGGTAGATGTTGACTACGTCTATGTTCAGGATGGAAATTCACCGACCGTGGCAAAACTGTTTCAGG<br>ATTATCATCTGTCTGAGGTGCTAAAACCCGATAAAATCGGGTTCTTCTTCGACCATTCAGTTCTGGTACCTGATAAAACCAT<br>GGCTAAACGTGTCAACGAGGCCATGGAATTTGCAAAAAAACTTGGAATAAACATCTATTCACGAGGGGAGGGAATTAGTCA<br>CGTCATTGCCCTGGAGAGTAAAATATTTAAACCCGGCAATATAGTGCTGGGCGCAGATTCCCATACTTGTACAGGGGGGG<br>CCGTACAGTCTTTAGCGCTGGGAATGGGGGCTACCCAGCTTTCTTGTACAAAGTGGT |
| **delhvrD** | ACAAGTTTGTACAAAAAAGCAGGCTAAGTTGCTATGGCATTGCACTTTTTCCACGGGGTTGATTTAGGGCTCGATTTAACAA<br>AATTAATCGCGTTGAGTGAAACGGTCGCCAGGTACAGCCATCAAAAAATCAGTCCATGGCAGCCGATCGTAGGCGATAAC<br>GTTTTTGCACATGAATCGGGCATTCACGCAAATGGTATGCTCAAAGACAGCAGTACTTTTGAACCCTTCGACCCAGCTACG<br>GTGGGAGGAGAACGACGTCTGGTCGTGGGTAAACATTCCGGTCGCGCCATTATCAAACATTTTCTCGAAGAATCAGGCGT<br>GAAAGCTGCCGACGATAAGGCTCTTGATCGCTGTTTAGAACGCGTGAGAAGTCATGCCGTGCGCCACCCCGGTGGGATC<br>CCTCCACATGTATTAGTTGATCTGTATACCGCGGGGTAAACAGGTATGAAAAGTAATCAGCCGCCCGGGGCATTGAACGG<br>CACTGTTCCACATTTTGAGGGAGAGTCCGCATATGAGTAAAACTGCTGAAAATTATCGCGTCAGACGCGTGGAAGGGAAC |

**Commented [BHK1]:** I feel bad I told you to use smaI. That's a unique enzyme for pDONR1K18ms but not for pR6KT2G

| | |
|---|---|
| | ATCTCGACCGACGATATTATACCTGCGCGCTATAAACATATGTATACCGAGCCGGCCCAGCTGGCACCGCATCTTTTTGAG<br>AGCCGTTTTCCCGGATTTAGGGAAACGCTCAGTATCAATGATGTGCTTGTATGTGATCAAATATTCGGTATAGGGAGTTCG<br>CGGGAGCAGGCAGTAACAACTCTGTTGGCATGTGGTGTTAAATATGTATTTTCCCCTTCTTTCGGGAGGATTTTTTTTAGGA<br>ACTCTTGGAATTTGGGTTTACATGCGATTGAGGTCGATACGAGTGAACTTGCAGATTTAAGTGAAATTAAAATAGAACTGAC<br>TGGAGGGGTAATTTATACAGAAAATAATCAAACCCAGCTTTCTTGTACAAAGTGGT |
| **delhvrE** | ACAAGTTTGTACAAAAAAGCAGGCTTATGCTAAACCTGCAAATGACTTTGAGGGTCTGAAATTTGATTATATTTTCATTGGA<br>AGCTGTACTAACAGCAGACTTGAAGATATCAAAGAGGTGGCCGAAATTGTTGCTGGTAAAACAATACATCCCGATATTCAC<br>TGCCTTCTGACGCCAGGATCGAAAAGTGTTTATCTAAAAGCTCTCCAGGCGGGATATATCGATACGCTTATCCGCTCGGG<br>CATTATTGTCACCCCACCGGGTTGTGGAGCTTGTGTGGGTACCCAAGGAACCATTCCTGCGGATGGGGAGAAAGTATTAA<br>GTACGATGAACCGCAATTTTAAGGGAAGAATGGGGAATGCTGAGGCAGACATCTTTTTATGTTCTCCACGAACTGCAGCGA<br>TGGTTGCATTGAACGGCACTGTTCCACATTTTGAGGGAGAGTCCGCATATGAGTAAAACTGCTGAAAATTATCCCGGGAAT<br>AAAATTATGGAAAAAAAAGGTGATATTTTAAGAGGTTATAGCAATGAAAAGTGAAAAGTTTGATGGTTTGGCTGATAACTAT<br>GATAAATATCGTCCCCGTTATCCTGCAATCCTTTTCAAGGAAATCCATGACTGGATGCAGCCGTCTGCCAAAAATATATACG<br>ATATTGGCGCAGGCACAGGTATTGCTATTGAAGGTATGACACGTGTCACTGGAAAACACTATGATTTCACGGCGATAGATA<br>TTTCTGAAGATATGATAAAAAAAGGAAGGGAAAAACTGCCTGGTACGACTTGGGTTAAAGGAAAAGCGGAAGATATTCTTT<br>CTGATAAAAGCCGTATTGACGTCATTATGGCGGCACAGTCCTTCCAATGGATGGATAGAGCTAAAACATTAGAAGTTTCGA<br>TAAAATCTTTAAATAAGGGCGGGGTTTTTGCAGTTTTGCAAACCCAGCTTTCTTGTACAAAGTGGT |
| **delhvrF** | ACAAGTTTGTACAAAAAAGCAGGCTTTTGAGAGCCGTTTTCCCGGATTTAGGGAAACGCTCAGTATCAATGATGTGCTTGT<br>ATGTGATCAAATATTCGGTATAGGGAGTTCGCGGGAGCAGGCAGTAACAACTCTGTTGGCATGTGGTGTTAAATATGTATT<br>TTCCCCTTCTTTCGGGAGGATTTTTTTTAGGAACTCTTGGAATTTGGGTTTACATGCGATTGAGGTCGATACGAGTGAACTT<br>GCAGATTTAAGTGAAATTAAAATAGAACTGACTGGAGGGGTAATTTATACAGAAAATAATCAAATAAATTTTTTCCCTCCCAG<br>CTCGCAGATGACGGCAATTGTCAGTGCAGGTGGCATAATACCCTACACCATAAATAAAATTATGGAAAAAAAAGGTGATAT<br>TTTAAGAGGTTATAGCAATGAAAAGTGAAAAGTTTGATGGTTTGGCTGATAACTATGATAAATATCGTCCCGGGATAAGTGA<br>GTTATTTATAGCTAAGAAGCGTGATGATTCATAGCATGACGATTCAGGACATCAGCATTGAACAAGCTTCGTATGAAGATG<br>CGAAGCTGTTAAGAAAGGCTTTAGAAAAAGTTTACGAACCTTATACACTGAATTTCTCACCTACCGCTTTGCAGTTTACTGA<br>AAATATCATTGCTCAGGAATCCTCGAAGTGGCTAGTCGCGAAATACAAGTCAGACATTGTGGGGGCAGTGAGATATGAACT<br>TTATGATATTTATCTGGACTTCCATTTTCTCTGCGTCACACCACCATTCAGAAAAATGGGAGTAGGGAATGAATTATTTCATA<br>AACTGAAGAAAATAGCGTATGAAAAAAGAAAGGATTTTATGAAGATCGTTTTGCGAGATTCGCTAAGCTATAACCGACGCTA<br>TTTTGAAAGTAAAGGATTTTACTTTTATCATAAAACCCAGCTTTCTTGTACAAAGTGGT |

| | |
|---|---|
| **delhvrG** | ACAAGTTTGTACAAAAAAGCAGGCTAAACATTAGAAGTTTCGATAAAATCTTTAAATAAGGGCGGGGTTTTTGCAGTTTTGC<br>AAAACAATCGAGATTACAGAAATAATGAAATGCTTAACAAATATGAAGGTTTGCTAGAGAAATTTAGTCCAGGTTACAGCAG<br>ACATTATCGCGACTATGACTATGAAAATGAAATCACCAATGTTTTTAAATTGCCTATTGCTAACTTTAAGAAAGTCGTCACAG<br>GGTGGACTATGGAAATGATTTCAGAAGATTTTTTTGGATTCATTTCTTCCTCAACCCAGGTACAAAGAGCTATTGAGAACGA<br>TCGTAATGGATTCTGGAAGGAGATTGAAATCTTGATTGACGAACACTCAGTTGGTGGAAAAATTAGCATAGATTATATAAGT<br>GAGTTATTTATAGCTAAGAAGCGTGATGATTCATAGCATGACGATTCAGGACATCAGCATTGAACAACCCGGGCATAGTGT<br>ATTTATTTTAAAATTAAACGGTGAAAAGCCATGAATAAAAAAGCATTGGTTATTGGGTTGAAAAGTAATATGGAAAGAGTCAT<br>CAAAGGGCTCAATGAGATAGAATTTATTATTATTGACAGAGGAACACTGGATAACGAAAGTATAGATTATATCATTAATCTTT<br>CAGATGATTTAATGCATAAAAAATCATTCAGTTATGTTATAGCCAGTTCTGAGGATTTTATCGCTTTGGCTGGGTTATTGCGT<br>AATCGTTATTCACTTTATGGTGAAAAATATTATAAAAGTACAATTGCAACCAATAAATTCCTGATGCGTAATTTCTGCTCAGG<br>TTTTTTATCCTGTCCAAAGTTCTGGCTATCAGGTGAGATCATCAATTCAGAGAATCTTTTACTGTCATCCCAGAAAGATTACA<br>TCGTAAAACCTCTTACAGGTAGTTCTGACCCAGCTTTCTTGTACAAAGTGGT |
| **delhvrH** | ACAAGTTTGTACAAAAAAGCAGGCTGCTGTTAAGAAAGGCTTTAGAAAAAGTTTACGAACCTTATACACTGAATTTCTCACC<br>TACCGCTTTGCAGTTTACTGAAAATATCATTGCTCAGGAATCCTCGAAGTGGCTAGTCGCGAAATACAAGTCAGACATTGT<br>GGGGGCAGTGAGATATGAACTTTATGATATTTATCTGGACTTCCATTTTCTCTGCGTCACACCACCATTCAGAAAAATGGGA<br>GTAGGGAATGAATTATTTCATAAACTGAAGAAAATAGCGTATGAAAAAAGAAAGGATTTTATGAAGATCGTTTTGCGAGATT<br>CGCTAAGCTATAACCGACGCTATTTTGAAAGTAAAGGATTTTACTTTTATCATAAATACCAGACAAATATGCATAGTGTATTT<br>ATTTTAAAATTAAACGGTGAAAAGCCATGAATAAAAAAGCATTGGTTATTGGGTTGAAAAGTAATCCCGGGTTGCATGATAT<br>TTCTGACAGACAAGAGTAAGATTCGATATGTTAAAATATGAAACCAAAAAAAACGCTTTCTTCCAGATATACAATTTTCTTTC<br>ATGCTCGGCTGAAGGGATTCAGACGGTTATATTTTTGTGGCTTATATATCATGAAACTCATAGTCCGATGCTTGTCAGTCTG<br>ACTATAGTTTCTTCATATCTTCCATCAGCCGTATTAGGATTCTTTTTCCTCAAAAAAGCTGATGCGAGCAGTCCTGGAAAAC<br>AGTTATTTATCAGTAATGTTTCTCTTAGTGCCATATCGCTAATTGTCTATTTTATTTTAATGAGAAATGAGGGTTTTGAACTAA<br>TCACGCTCAGTATCTTTTATTTGGCACAGGCCGTATTATCTGTTGTCAAAATGTTTAATAAGACATCCCAAAATCGCATTATC<br>AGAACAGCATTCAGTAATAGTGATGACCCAGCTTTCTTGTACAAAGTGGT |

| | |
|---|---|
| **delhvrI** | ACAAGTTTGTACAAAAAAGCAGGCTGAAATGACTAATGGCGTTTTTCATATTGAAATGTATCATACTCAGGATGGATTTATA<br>CTCGGTGAGTTTGGTATCCGCCCTCCCGGAGCGGGTGTGACCGATATGTATTATATGTATAGAGGCGTCGATTTCTGGGA<br>GGCATTTATTTATTCTCAGATTGATAAAGAATTTATTTTACCCCTAAATCATAAGTCCGATAAATATTGTGCTGCTATTGGGA<br>TATGTTCTTCTTTCCCGGTCGATGATATCAGAAGTACGTCAAAAGTATCCGTGGATAAATATGTAAATTTGCGTGAGAATGC<br>TGCTAAACCCGCAGTTCCCAGTTCAACGTCTTTTAATCATATGATTTATGTTTCTTCATCATCCTTAGAAGAAATAAGGAATT<br>TTTTGCATGATATTTCTGACAGACAAGAGTAAGATTCGATATGTTAAAATATGAAACCCCCGGGGTTTATCAGAGTCGCAGT<br>TTTAACTATGAGGATTTAGGTGATGTTAAATTTCAAAAGTGATTTTTCTGAATTTATTACTGGTTTTTATCTTAAACAGTTTAC<br>TCATTTAAATACGCAGGAAAGAGAACATGTGCTTGAAACGCTTGGCGTCACACCCAGCGCAATCAATGAGTTTATAACATC<br>AGAAGATATTTATATTACCTTACCTCATGCAAGTATGAATGTTTTTTTTCCAAGAGCGGGGGTCTCACGTTATGTTTACGATC<br>TACAAAATAGTAACAAAAACGCGTTCCATTTAAGGTTTTTCTTGACCCATACCAACTTTAGCGATCTGAACTGGCGCCCTTA<br>TGCATGGTGGTTTAATAACGGCGGGAAAATAGATAAACTTACCTTTTTCACACGAAATACCCAGCTTTCTTGTACAAAGTGG<br>T |
| **delhvrJ** | ACAAGTTTGTACAAAAAAGCAGGCTGCAGCACTCAATATGCTTTCTCTAAAGTCTTCATTACGGTTTTCTCTGTGTATTTCTG<br>GCATCTGTCTGATTTTGATGTGTTTATCAACGAGAAATTATTATGCCGTATACTTTTTTTTATTTCTCACTTCTTTTTTTGTTTC<br>ATGGCATATGATTTCTATTAAGGTACTCACGAATCAAATGCCTGATATAGAAAATATTGGTAAATTCACAATGATGAGAAACT<br>CAGTCGCTTCTGGTGTGAAAATTGTCTTTTCTATTTCGTCTGGTGCTTTTTTGACTTTTTACAGCATAACTACAACTTATCTA<br>ATACTTGCGATACTGCTTATTTTTTTCAATGTATTATGGGTTTATCAGAGTCGCAGTTTTAACTATGAGGATTTAGGTGATGT<br>TAAATTTCAAAAGTGATTTTTCTGAATTTATTACTGGTTTTTATCTTAAACAGTTTCCCGGGCAGGAACATGATCTTTTTTCAA<br>AGAAAGAATAATAAAACATAAACTAGAAATAACCTCTCATTTTGAAAAGGCGACATGGTTACCGGGTTGAAGGTTGTATATA<br>AGAACCAGTTTAAAATATTTTAAAGTTACATGTCATTGCTCATTTAACATCATGCGATAACACCTCAACACTCATTGGAGCTC<br>TATTATGCTAGATAAAAGTGCTTTTCGCCACGCAATGTCGCATCTTCCTACTGCAGTCACTATTGTAACATCAAGCGGCTCC<br>TGCGGGGCTGCAGCATGTACAGTTTCATCCGTGTGTTCTGTCACCGACGACCCACCCACTTTACTCGTATGTATAAATCGT<br>GCCTCCAATAACAATAGTGTTATAAGAAATAATGGCTCTTTGTGTGTCAGCATTCTTTCAGGAGAGCAAAGCAACATTGCAA<br>TGCAATGTGCTAATCATACCCAGCTTTCTTGTACAAAGTGGT |
| **delhvrK** | ACAAGTTTGTACAAAAAAGCAGGCTAATTTTACAAACATTGCGCTGTTAAATTTTTTTCAGCCTGCAGCATTTGTGGGAGGC<br>GAAAAAATGGATAACTACTGGCATCAGGTGATTGAAAAATGGAAAATGGCATTACCAAATCAATCTGAAGTTGAGTTTGCGT<br>TACCCACTAATCTGGTCATGCCTGCCGTTCAAGAGTATATTTACCCTTATAAACCATCAAGCGATATAGCAGAACAGCTTAT<br>TAAAAATAACATCCCATACAGTCTTACCATGGCTATTCAGGAACATGATCTTTTTTCAAAGAAAGAATAATAAAACATAAACT<br>AGAAATAACCTCTCATTTTGAAAAGGCGACATGGTTACCGGGTTGAAGGTTGTATATAAGAACCAGTTTAAAATATTTTAAA<br>GTTACATGTCATTGCTCATTTAACATCATGCGATAACACCTCAACACTCATTGGAGCTCTATTCCCGGGGGCACTATGCCTA<br>AAATGCTGTAATACTGTCTTTATTTCGTGGCATAAATATTCTGAGAAAAAACAGGCCGCATCATATTCATCAGATACAATCAC<br>CAGGTGTTAATATTCTTGTTTCTACATTAAAATAGAGTTGTAAGGTGTGTTTTTTTTAGGTGTTTAAATTATACCTTCCGTGTA |

| | |
|---|---|
| | AATATTAATTTTTTATTTGATTTTCACTTTCATTTAAATTCTGAATTTTTGCTTCCTTTTTCCTGAACCCATTATACCTGAATCC<br>CAATTTAAATCAGATTCGGGAACCAGGTCAGGAAGTCAGGAAGGCCGTCAGCAGGAATCAGGGGATCTTCATCAACTATG<br>TGTCCGTAATTAAGACGAAGGTTAAGCTGAGTTTTTACGTAATGCACTATAATTTTTGCAGAAGCCGTCTAATCATAATTCA<br>GTAACATAAGAGGATTAGTCATCACCCAGCTTTCTTGTACAAAGTGGT |
| **Pre-SNARE** | atgtcaactgggttcgtgcgagctcatcgatgcatacaagtttgtacaaaaaagcaggctataaaaaaacgcccggcggcaaccgagcgttctgaacggtgcctaatgagtgagct<br>aactcacattaattgcgttgcgcTCATTGACCGGACTCAAGGCGCGAAACTTGACGCGCAAGTTGCATAAGAGAATCGGCTAACGCGC<br>GAGGTGATGCAGTCTGAGTATTGGGTGCCAGAGTTGTTTTGCGCTTAACCAGGGATACGGGTAACAACTGATTTCCCTTCA<br>CCGCCTGACCTTGCGAAAGTTGAAGCAGGCGGTCTACGCTGGTTTGACCTAACAAACGAAAATCTTGTTTGATTGTAGTCA<br>GCGGAGGAATGTAACACGAAGAGTCCTCGGTGTCATCATATCCGACGACAGAGATGTCGGCCCCAACACGAAGTCCCGA<br>CTCTGTGATGGCACGCATTGCGCCCAGTGCCATCTGGTCATTGGCTACCAGCATCGCCGTGGGAACAATCCCCTCGTTCA<br>GCATCTGCATTGTCTGCTGAAAGCCGCTCATAGCTGACCAGTCACCCTCACGTTCGGCAATCGGTTGAATTTGATTGCGA<br>GTCAAATACTTGTGCCAGCCAGCTAAACGCAGGCGAGCTGACACGCTGCTAAGTGGCCCCGCAAGTAAGGCAATTTGTTG<br>ATGGCCTAAAGCAACCAGATGTTCAACGCCCAGACGAGTACCATCCTCGTGCGAGAAGATAATACTGTTAATGGGTGTTTG<br>ATCGCTTACATCCAGAAACAGGGCGGGTACGTTCGTACAAGCTGCCTCCACCGCGATTGCGTCCTGATCATCCAACGGGT<br>AATTGATGATTAAGCCACTCACACGCTGAGCCAGTAAGTTATGCACGGCTGCTTTACAAGCCTCCACACCGCTACGTTCAA<br>CCATGCTCACTACAACTGATGCACCCAACTGGTCTGCGCGGGACTTGATAGCCGCGACGATTTGACTAGGCGCATGAAGT<br>GCCAAGCTTGAGGTCGCGACTCCAATAAGCAAAGACTGCTTGCCGGCAAGTTGTTGGGCGACACGGTTCGGGATATAATT<br>CAATTCAGCCATTGCTGCCTCAACCTTCTCGCGTGTCTTAGCGGAGACATGAGATGCCTGGTTAACCACACGCGATACTGT<br>TTGATATGAGACCCCCGCGTACTCTGCGACGTCATATAATGTCACTGGTTTCACattcaccaccctgaattgactctcttccgggcgctatcatg<br>ccataccgcgaaaggttttgcaccattcgatggtgtcacggttctggcaaatattctgaaatgagctgttgacaattaatcatcggctcgtataatgtgtggaattgtgagcggataaca<br>atttcacacaggaaacagaattgatccggccaagcttgtaagaaggagatatacatatgatcaaaaaacttattgcagaaaagggtactctgatttttattgaggcccataatccgct<br>ctccgcattaattgcgtctaaagcagaacaaactaattcagaaggccgtattgtcaaatttgacggtatatggtcaagctcgttaacggactcagcatctcgcggtattcccgataacg<br>aaacactggcattaagcagcaggttagaaaatattgctgatatccgaaatgtgacagacatgcccatcatcatggatgctgatacgggggaaaaccagaacattttagttattacg<br>taaaaagaatgattaacaacggtgtaaatggcgtcatcatcgaagataaaacaggattaaagaaaaattctttgttcggcactgaagtagaacagactctcgTctagaggtaccG<br>gcgcgccagactcgctagcgcggccgcactagtaggcctacccagctttcttgtacaaagtggtgcatgcgatatcgagctctcccgggaattccacaa |

**attB* sites are highlighted in yellow and green highlight indicates restriction site

**Table S3.** A list of primers used for cloning and sequencing in this study

| Primer | Primer sequence (5'-3') | Annealig temp. (ºC) |
|---|---|---|
| **Out primers for deletion mutation confirmation** | | |
| Out_hvrB_F | AGTAGGACAGACTCTCGCAGA | 58 |
| Out_hvrB_R | GCAACTACCTGTGCATACTCCT | |
| Out_hvrC_F | GGGCGAATGGAAAAATACGGA | 55 |
| Out_hvrC_R | AGAAATGGTGTTTGTCCGTACT | |
| Out_hvrD_F | ATTGAAGCAGGAGCACGCTA | 58 |
| Out_hvrD_R | TGGTGTAGGGTATTATGCCACC | |
| Out_hvrE_F | TCAGTGGCAGACAGTGTCATC | 58 |
| Out_hvrE_R | AGTCGCGATAATGTCTGCTGT | |
| Out_hvrF_F | CGTGGAAGGGAACATCTCGA | 58 |
| Out_hvrF_R | ACTTTTCAACCCAATAACCAATGCT | |
| Out_hvrG_F | ACTTGGGTTAAAGGAAAAGCGG | 58 |
| Out_hvrG_R | TCATCTCTCATTAAAACCTTTTCTTCA | |
| Out_hvrH_F | GTGAGTTATTTATAGCTAAGAAG | 50 |
| Out_hvrH_R | GCATGAGCGCTTGCATGTAG | |
| Out_hvrI_F | CGACAGGCCTATACTCGAGG | 60 |
| Out_hvrI_R | CGCAGACGTCGATCTACAGA | |
| Out_hvrJ_F | ACACATCCCTTCCTTCATTATCT | 55 |
| Out_hvrJ_R | CAACTACAGCAGGAGACCC | |
| Out_hvrK_F | TGCAGAAGACTGAATGGCGA | 58 |
| Out_hvrK_R | TGTGCCTGATTCTCACTTCAA | |
| **Comp primers for complementation** | | |
| Comp_hvrB_F | ggggacaagtttgtacaaaaaagcaggctTCTTGCATGAGTGTAAAGGAA | 50 |
| Comp_hvrB_R | ggggaccactttgtacaagaaagctgggtCTCGCAATACATTATTGATT | |
| Comp_hvrC_F | ggggacaagtttgtacaaaaaagcaggctACTGGGGCTTAAAAGACCTACT | 55 |
| Comp_hvrC_R | ggggaccactttgtacaagaaagctgggtTTACCCCGCGGTATACAGATCA | |
| Comp_hvrD_F | ggggacaagtttgtacaaaaaagcaggctTGATCTGTATACCGCGGGGTA | 55 |
| Comp_hvrD_R | ggggaccactttgtacaagaaagctgggtGACGCGATAATTTTCAGCAGTT | |
| Comp_hvrE_F | ggggacaagtttgtacaaaaaagcaggctATGGTTGCATTGAACGGCACT | 55 |
| Comp_hvrE_R | ggggaccactttgtacaagaaagctgggtCAGCCAAACCATCAAACTTTTC | |

| | | |
|---|---|---|
| Comp_hvrF_F | ggggacaagtttgtacaaaaaagcaggctTGGCATAATACCCTACACCATA | 55 |
| Comp_hvrF_R | ggggaccactttgtacaagaaagctgggtCTATGAATCATCACGCTTCTTAG | |
| Comp_hvrG_F | ggggacaagtttgtacaaaaaagcaggctGTGAGTTATTTATAGCTAAGAAG | 50 |
| Comp_hvrG_R | ggggaccactttgtacaagaaagctgggtCCTTTGATGACTCTTTCCATA | |
| Comp_hvrH_F | ggggacaagtttgtacaaaaaagcaggctACCAGACAAATATGCATAGTGTATT | 55 |
| Comp_hvrH_R | ggggaccactttgtacaagaaagctgggtGAATCCCTTCAGCCGAGCAT | |
| Comp_hvrI_F | ggggacaagtttgtacaaaaaagcaggctTCTGACAGACAAGAGTAAGAT | 50 |
| Comp_hvrI_R | ggggaccactttgtacaagaaagctgggtAAGCGTTTCAAGCACATGTT | |
| Comp_hvrJ_F | ggggacaagtttgtacaaaaaagcaggctAGTCGCAGTTTTAACTATGAG | 50 |
| Comp_hvrJ_R | ggggaccactttgtacaagaaagctgggtGAGAGGTTATTTCTAGTTTATG | |
| Comp_hvrK_F | ggggacaagtttgtacaaaaaagcaggctAACACCTCAACACTCATTGG | 53 |
| Comp_hvrK_R | ggggaccactttgtacaagaaagctgggtTGCCTTATGATACACGATGAT | |
| **pR6KT2G-specific primers** | | |
| pR6KT2G_F | GTCTTAAGCTCGGGCCCC | 60 |
| pR6KT2G_R | GGGATATCAGCTGGATGGC | |
| **Pre-SNARE amplification primers** | | |
| Pre-SNARE_SphI_F | ttctagaggtaccgcatgATGTCAACTGGGTTCGTGCG | 58 |
| Pre-SNARE_NsiI_R | gcagaaagattaaatgcaTTGTGGAATTCCCGGGAGAG | |
| **pSNARE-specific primers** | | |
| lacIQ_F | TCGATGGTGTCACGGTTCTG | 55 |
| test_pSC201_R | CACATGTGGAATTGTGAGCG | |
| **Sequencing primers** | | |
| M13_F | GTAAAACGACGGCCAGT | 60 |
| M13_R | CAGGAAACAGCTATGAC | |
| hvrA_F | CTCATCTTTACGCATCAAAGGTG | 58 |
| hvrA_R | TGTTCCATACGATATATAGGCAC | |
| hvrI_F | TCTGACAGACAAGAGTAAGAT | 55 |
| hvrI_R | AAGCGTTTCAAGCACATGTT | |
| **Allelic exchange primers** | | |

| | | |
|---|---|---|
| hvrA_exc_F | GGGGACAAGTTTGTACAAAAAAGCAGGCTTTAGGCAGG CACTCTATAAGAGC | 58 |
| hvrA_exc_R | GGGGACCACTTTGTACAAGAAAGCTGGGTTAACCTGCC ATGCTTCCCAG | |
| hvrB_ex_F | GGGGACAAGTTTGTACAAAAAAGCAGGCTGCAACCAGC AACAGAGAACTC | 55 |
| hvrB_ex_R | GGGGACCACTTTGTACAAGAAAGCTGGGTTCCGTATTTT TCCATTCGCCC | |
